# A multi-input optic glomerulus mediates opposing behavioral responses to visual objects

**DOI:** 10.64898/2026.01.26.701771

**Authors:** Inês M.A. Ribeiro, Wei-Qi Chen, Nikolas Drummond, Stefan Prech, Michael Sauter, Alexander Borst

## Abstract

Prey, predators or conspecifics are first detected as visual objects in many seeing animals. Vision guides behavioral actions towards or away from these objects. An error in this visual perception could prove fatal. How object information is untangled to avoid errors remains unclear. Here we show that LC10d visual projection neurons in *Drosophila melanogaster* mediate avoidance of visual objects in the absence of a chemosensory profile. LC10d neurons are broadly tuned to objects and project to the same retinorecipient brain region that receives inputs from LC10a neurons, which are required for tracking. The descending neurons DNa10 are directly downstream of the anterior-facing LC10d sub-population and mediate LC10d-dependent avoidance. Our work demonstrates the use of two similar neuron types and chunking of the visual field into zones as strategies to disentangle similar sets of visual cues requiring nearly opposite behavioral responses.

## Introduction

Spotting a predator, a competitor or a future mate at a distance likely starts with the detection of a small, ambiguous object in many seeing animals. The enveloping sensory context, composed of odors, tastants and sounds, influences whether the object is interpreted as harmless or an actual threat (Budick and O’Malley, 2000; Land and Collett, 1974; Nikonov et al., 2006; Nordstrom et al., 2006). The absence of a chemosensory context or tell-tale cues suggestive of an imminent attack, such as looming, together elicit avoidance. In contrast to an outright escape, avoidance may precede attraction at a later time point if the appropriate sensory context arises (Cheng et al., 2019; Maimon et al., 2008). How ambiguous sets of visual cues, that could equally represent a mate or a competitor, are resolved in central neural circuits remains unclear. Despite its low spatial resolution, vision is a major sensory modality in most insects, with high temporal acuity and large portions of neural real estate dedicated to processing visual input (Strausfeld, 2012). Furthermore, vision affects several aerial and terrestrial behaviors in insects (e.g., (Agrawal et al., 2014; Briceno, 2003; Land and Collett, 1974; Li et al., 2024; Nordström et al., 2008; Olberg, 2012; Ribeiro et al., 2018; Thyselius et al., 2018).

At the level of photoreceptors, visual input starts as an abstract percept that is processed in the optic lobe in invertebrates, where visual features, like color or motion, are extracted (Borst and Groschner, 2023). Visual projection neurons relay processed information to the central brain, forming a bottleneck between the optic lobe and central neural circuits regulating behavior (Cheong et al., 2020). In *Drosophila melanogaster*, lobula columnar (LC) neurons are visual projection neurons composed of many cells that tile the lobula neuropile in the optic lobe and collectively cover the visual field (Fischbach and Dittrich, 1989; Nern et al., 2025; Otsuna and Ito, 2006; Wu et al., 2016). LC neurons relay combinations of visual features to the central brain that in some cases carry ethological meaning. For instance, LC4, LC6 and LC16 neurons all sense looming (Ache et al., 2019; Klapoetke et al., 2022; Wu et al., 2016). However, activation of LC4 or LC6 elicits a jump, whereas activation of LC16 leads to retreat. This suggests that, despite similar sensitivities, only LC4 and LC6 are associated with fast approaching threat.

Several distinct LC neuron types detect discrete objects, with varying degrees of specificity for size, contrast and speed (Keleş and Frye, 2017; Klapoetke et al., 2022; Ribeiro et al., 2018; Schretter et al., 2025; Städele et al., 2020; Wu et al., 2016). This apparent redundancy might boost flexibility in behavioral responses to visual objects. The circuits downstream different object-detecting neurons might provide a neural substrate upon which similar visual information is integrated with different multimodal sensory cues, such as chemical signatures. In this way, visually similar objects with disparate chemical signatures that merit different responses, can lead to different behavioral outcomes. Visual projection neurons might encode different features of the same object that are then combined to provide an enriched representation, thereby supporting the appropriate behavioral response, as was demonstrated for looming detection by LPLC2 and LC4 neurons (Ache et al., 2019). LC4 neurons encode expansion velocity and LPLC2 neurons encode large angular sizes of looming, converging onto the giant fiber to elicit escape (Ache et al., 2019; Von Reyn et al., 2017, 2014). Finally, different object detecting neuron types could be targets of various neuromodulatory pathways acting under different internal states, such as hunger or aggression, with responses to visual objects regulated in a top-down fashion (Alekseyenko et al., 2019; Bertsch et al., 2025; Hindmarsh Sten et al., 2021; Kohatsu and Yamamoto, 2015; Ribeiro et al., 2018; Rubin et al., 2025; Schretter et al., 2025). Whether one or more of these circuit mechanisms are at play in object detecting projection neurons in Drosophila remains incompletely understood.

Among behaviors involving object detection, social interactions stand out. Maintaining close proximity to the female is a pre-requisite for a successful courtship and eventual copulation. During courtship, the male orients towards and chases the female, while singing a species-specific song (Hall, 1994; Rings and Goodwin, 2019; Yamamoto and Koganezawa, 2013). Tracking the female relies on vision and depends on LC10a, and, to a lesser extent, on LC9 neurons (Agrawal et al., 2014; Bidaye et al., 2020; Cook, 1980, 1979; Cowley et al., 2024; Markow, 1987; Nojima et al., 2021; Ribeiro et al., 2018). LC10a neurons are tuned to moving objects with angular sizes and speeds within the range experienced by a courting male (Ribeiro et al., 2018), and project to the anterior optic tubercle (AOTU) (Otsuna and Ito, 2006), the largest optic glomerulus in the fly brain. At least six other neuron types from the LC10-group project to the AOTU central unit, most displaying a spatial arrangement of their terminals that represents the fly visual field with coarse retinotopy (Dorkenwald et al., 2024; Hoeller et al., 2025; Nern et al., 2025; Plaza et al., 2022; Schlegel et al., 2024; Wu et al., 2016; Zheng et al., 2018). LC10d and LC10bc are dispensable for maintaining close proximity to the female during male courtship behavior (Cowley et al., 2024; Ribeiro et al., 2018). Connectomic data reveal the existence of multiple parallel pathways stemming from the AOTU central unit, that combine information from two to three LC10-group neuron types with varying degrees of inter-connectivity (Berg et al., 2025; Dorkenwald et al., 2024; Plaza et al., 2022; Scheffer et al., 2020; Schlegel et al., 2024; Zheng et al., 2018). The map-like spatial organization of the AOTU-incoming visual inputs suggests that AOTU-output neurons might synthesize spatial information to encode object position or speed, or even predict future positions. Alternatively, the AOTU spatial information could be summarized relative to specific zones in the fly visual field, such as anterior or posterior, dorsal or ventral.

Here we reveal that inputs to the AOTU regulate at least two opposing behavioral responses to visual objects, tracking and avoidance. Single pair social interactions were tested in paradigms that expose the observer fly to an expanded visual feature space, leading to the occurrence of visual object avoidance-like behavior before courtship is initiated. We show that LC10d neurons, but not LC10a, are required for visual object avoidance in different behavioral paradigms, including vision-only and fly-on-ball assays. LC10d neurons are sensitive to discrete visual objects, and are overall more broadly tuned than LC10a. The descending neurons DNa10 receive input from a sub-population of LC10d cells representing the anterior visual field (Dorkenwald et al., 2024; Namiki et al., 2018; Schlegel et al., 2024; Zheng et al., 2018), and are necessary and sufficient for visual object avoidance. Our work showcases LC10a and LC10d as object detectors with overlapping tuning properties that mediate opposing behaviors. Moreover, we bring to light chunking of the anterior zone of the visual field as a mechanism of summarizing spatial information provided by the AOTU central unit.

## Results

### Visual object avoidance occurs in the pre-courtship period

In single pair assays, both the naïve male and virgin female are exposed to naturalistic sensory cues originating from the other fly. The major chemosensory cue that arouses the male to court, the contact pheromone 7,11-HD (Lu et al., 2012; Thistle et al., 2012; Toda et al., 2012), is detected when the male is in close proximity to the female, within tapping distance (Clowney et al., 2015; Koh et al., 2014). Odors influencing the male’s decision to court are detected at inter-fly distances below 5 mm (Clowney et al., 2015; Dweck et al., 2015). Visual objects at larger distances however are perceived without such chemosensory cues. In barrierless arenas with a 42-mm diameter, the first wing extension and following (or chase) bouts occurred roughly one minute after placing the male and female in the arena. For one third of the pre-courtship period, the female was perceived with an angular size below 10° (33.72% ±4.48s.e.m., N=32 pairs wild type Canton S), known to elicit visual object avoidance (Cheng et al., 2019; Maimon et al., 2008). The distribution of female positions relative to the male in heat maps of male-centered coordinates was frequently skewed to the posterior visual field during pre-courtship (Figure 1A, B). In contrast, female relative positions are highly enriched in a narrow, frontal visual field during courtship (Figure 1C). This suggests that male behaviour involves avoidance of the female before courtship is initiated, and then shifts to tracking during courtship.

**Figure 1:**
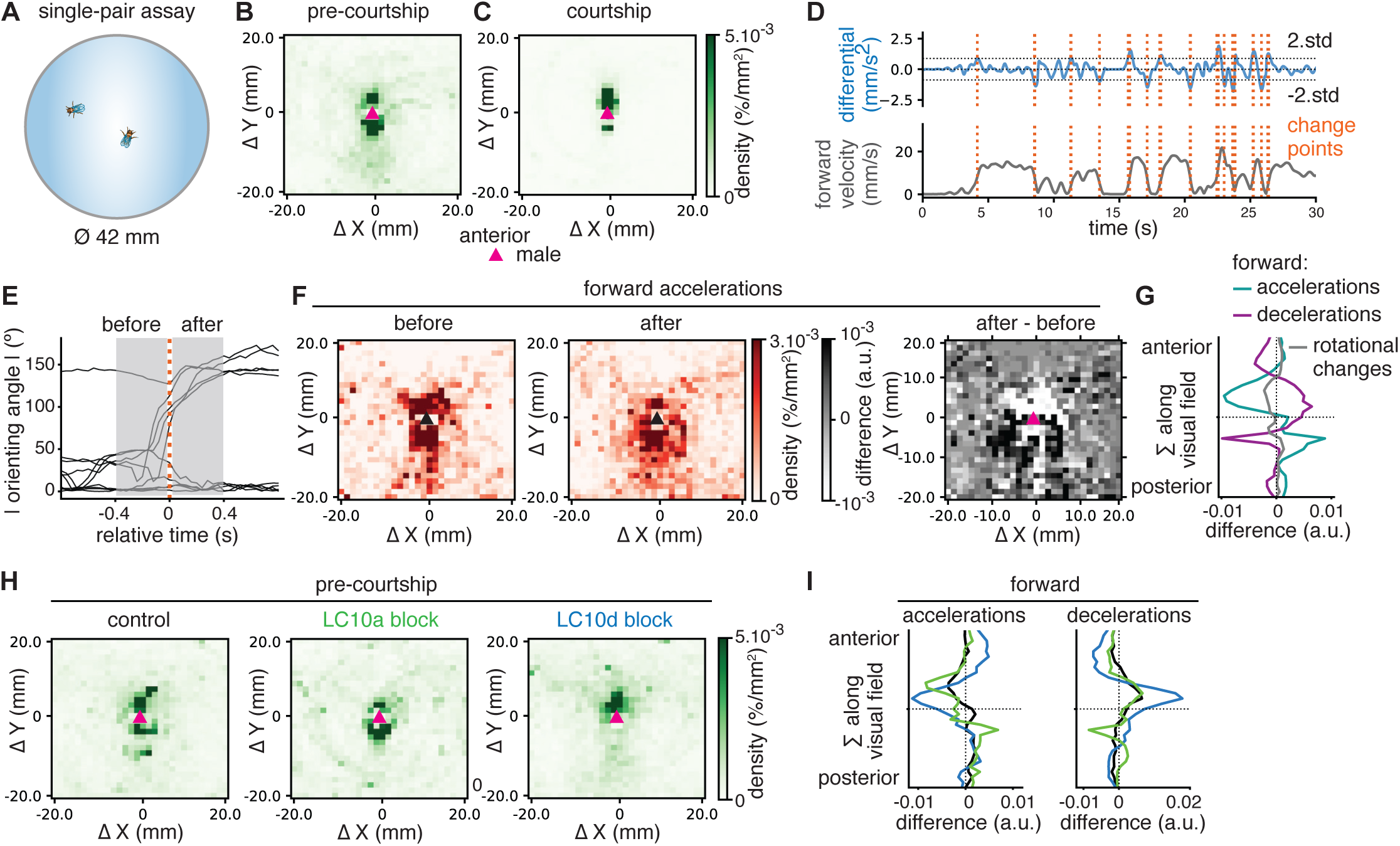
Visual object avoidance in the pre-courtship period in single pair assays. **A.** Schematic for single pair assays in arenas with a 42mm diameter. **B-C.** Male-centered heatmaps of female relative positions for the pre-courtship (B) and courtship (C) periods for Canton S single pair assays (N=32). **D.** Example 30 seconds trace of the male forward velocity (bottom, grey) and the first derivative (top, blue), with change points (dotted line, orange) overlayed. Local maxima correspond to accelerations and local minima correspond to decelerations. **E.** Absolute orienting angle for a set of aligned change points to indicate the amount of time before and after each change point was used to compute the difference heatmaps shown in F-G. **F.** Male centered heatmaps of female relative positions for 400 ms before (left) and 400 ms after (middle) accelerations in forward velocity, and the difference after – before map (right). **G.** Projection along the Y axis of the heat map after – before for forward accelerations (green), forward decelerations (purple), and rotational change points (grey), indicating the most likely outcome of female relative position along the anterior-posterior axis of the male visual field when the male changes his locomotion. **H.** Male-centered heatmaps of female relative positions for the pre-courtship period for single pair interactions for parental control (N=53), LC10a-SS1 block (N=14), and LC10d-SS3822 block (N=31). **I.** Projection along the Y axis of the heat map after – before for forward accelerations (left panel) and decelerations (right panel), for controls (black), LC10a-SS1 block (green) and LC10d-SS3822 block (blue). Mean area under the curve (A.U.C.) for the anterior visual field for accelerations with p > 0.05 for control vs LC10a or control vsLC10d block; for decelerations, with p> 0.05 for control vs LC10a block and p = 0.04 for control vs LC10d block, Student T-test.

To test whether males courting in 42-mm arenas rely on similar visual pathways reported previously using other courtship paradigms (Bidaye et al., 2020; Cowley et al., 2024; McKinney and Ben-Shahar, 2019; Ribeiro et al., 2018), we leveraged split-GAL4 lines targeting specific neuron types to express the neuronal silencer tetanus toxin light chain (TNT), thereby blocking neurotransmission in chemical synapses (Ribeiro et al., 2018; Sweeney et al., 1995; Wu et al., 2016). Median inter-fly distances and absolute orienting angles were significantly increased if LC10a or all LC10-group neuron types were silenced (Figure S1A-C). The proportion of time spent with wing extension was significantly reduced only upon LC10a-SS1 or LC10s-SS2 silencing, whereas following was significantly reduced upon silencing of LC10a-SS1, LC10s-SS2 and LC9-SS2651 neurons (Figure S1D,E). Courtship in 42-mm arenas thus involves female tracking with low values for absolute orienting angles and distances, and relies heavily on LC10a and LC9 neurons, as previously described (Bidaye et al., 2020; Cowley et al., 2024; Ribeiro et al., 2018).

In contrast to object tracking, visual avoidance does not require spatial precision. If the visual object is mobile, like the female, several locomotion maneuvers from both flies could result in a shift of the object away from the anterior field of the observer. To determine whether behavioral maneuvers carried out by the male result in visual object avoidance during pre-courtship, we used an unbiased analysis of locomotion. Male forward and rotational velocities were subject to a change point algorithm that extracted the timepoints in which the male modulated his speed (Figure 1D) (Wiltschko et al., 2015). The change points were aligned, and male centroid-centered heat maps of female relative positions were generated for 400 ms before and after the change point (Figure 1E, F). Female relative positions resulting from male-driven change in locomotion were obtained by subtracting the heat map corresponding to ‘before’ from the heat map corresponding to ‘after’ the change point (difference = after – before). Forward accelerations reduced, and decelerations increased, the likelihood of female relative locations anterior to the male, whereas large changes in rotational speed had little impact on female relative position distribution across the anterior-posterior visual field of the male (Figure 1G). All changes in locomotion combined resulted in an overall avoidance during the pre-courtship period, which was abolished when courtship was initiated and was absent in blind males and for randomly chosen points (Figure S1F-I). We conclude that the male actively avoids the female during the pre-courtship period, likely due to a disconnect between visual and chemosensory cues, and mostly employs forward accelerations accompanied by mild turns (Figure S2A,B) in the process.

### LC10d neurons mediate visual object avoidance during pre-courtship

LC10a and LC10d neurons co-project to the AOTU together with other LC10-group neuron types. Unlike LC10a, LC10d neurons are dispensable for directed courtship (Figure S1A-E)) (Cowley et al., 2024; Ribeiro et al., 2018). During the pre-courtship period, female relative positions were highly enriched in the anterior visual field for LC10d silenced males, compared to more evenly distributed anterior and posterior locations in controls and with a bias towards positions in the posterior visual field for LC10a-block males (Figure 1H). Examination of purposeful male locomotion maneuvers revealed that forward accelerations and decelerations increased the likelihood of frontal female relative positions upon LC10d silencing, compared to controls, with enrichment at high and low distances, respectively (Figure 1I, S1J,K). Large changes in rotational velocity showed a limited impact on female relative positions very close to the male in wild type, controls, LC10a- and LC10d-block males. Importantly, change point amplitudes in forward and rotational velocities were unaltered with genotype, whereas temporal profiles around change points showed negligible differences across genotypes (Figure S1L,M). Moreover, silencing LC10d neurons led to an increase in male – male interactions, which are typically very limited in the absence of food (Figure S1N,O) (Duistermars et al., 2018; Vrontou et al., 2006), suggesting that LC10d neurons coordinate avoidance in different social contexts.

To investigate whether changes in male locomotion during the pre-courtship period occur in response to specific female behaviors, female locomotion was examined in the time envelope centered on male change points. Female rotational speed was largely flat around male forward accelerations, decelerations and changes in rotational speed, whereas female forward speed showed small changes in the same time envelopes (Figure S2A,B, and not shown). Male forward accelerations were invariably accompanied by a mild turn in all genotypes examined (Figure S2B), which could underlie the alterations in orienting angle that we observed. The derivative of the absolute orienting angle male to female showed a nadir coincident with forward acceleration point, that was much deeper upon LC10d silencing (Figure S2C,D). The converse orienting angle, female to male, remained largely unchanged in the time envelope surrounding forward accelerations, decelerations and rotational changes, suggesting that the resulting change in female azimuthal position relative to the male is driven solely by changes in male behavior. In contrast to orienting angle, changes in the distance between the head of the observer fly and the tip of the abdomen of the other fly (distanceHT) were similar for male and female, showing a slight increase after male forward accelerations for wild type, control and LC10a block males (Figure S2E,F). The temporal profile of distanceHT differed between male and female only for LC10d block males. Analysis of changes in female locomotion and orienting in the time envelope around changes in male locomotion thus suggests that male-driven locomotion underlies avoidance.

We adapted a behavioral paradigm developed by Pan and colleagues, to assess male behavior in response to visual information from the female that is permanently disjointed from auditory, mechanosensory and chemosensory cues (Pan et al., 2012). In our 2-floors assay, a naïve male is placed alone in an arena stacked above another arena containing one virgin female. The flies are video recorded for ten minutes (Figure 2A). Female pursuit was absent in 2-floor assays for wild type or blind males (Figure S2G-J). Male centered heat maps of female relative positions for the entire 10-min assay showed a distribution to the anterior and posterior male visual fields, slightly enriched in the former (Figure 2B). This tendency to anterior locations relative to the male was absent when the male accelerated or decelerated (Figure S2G,K). Instead, modulation of male forward velocity resulted in a decrease of the likelihood of female relative locations in close proximity to the male, that was abolished in blind males and absent in randomly chosen points (Figure S2G-L). The 2-floor assay thus offers a paradigm to study behavioral responses to visual information from the female isolated from other sensory channels, with high throughput.

**Figure 2.**
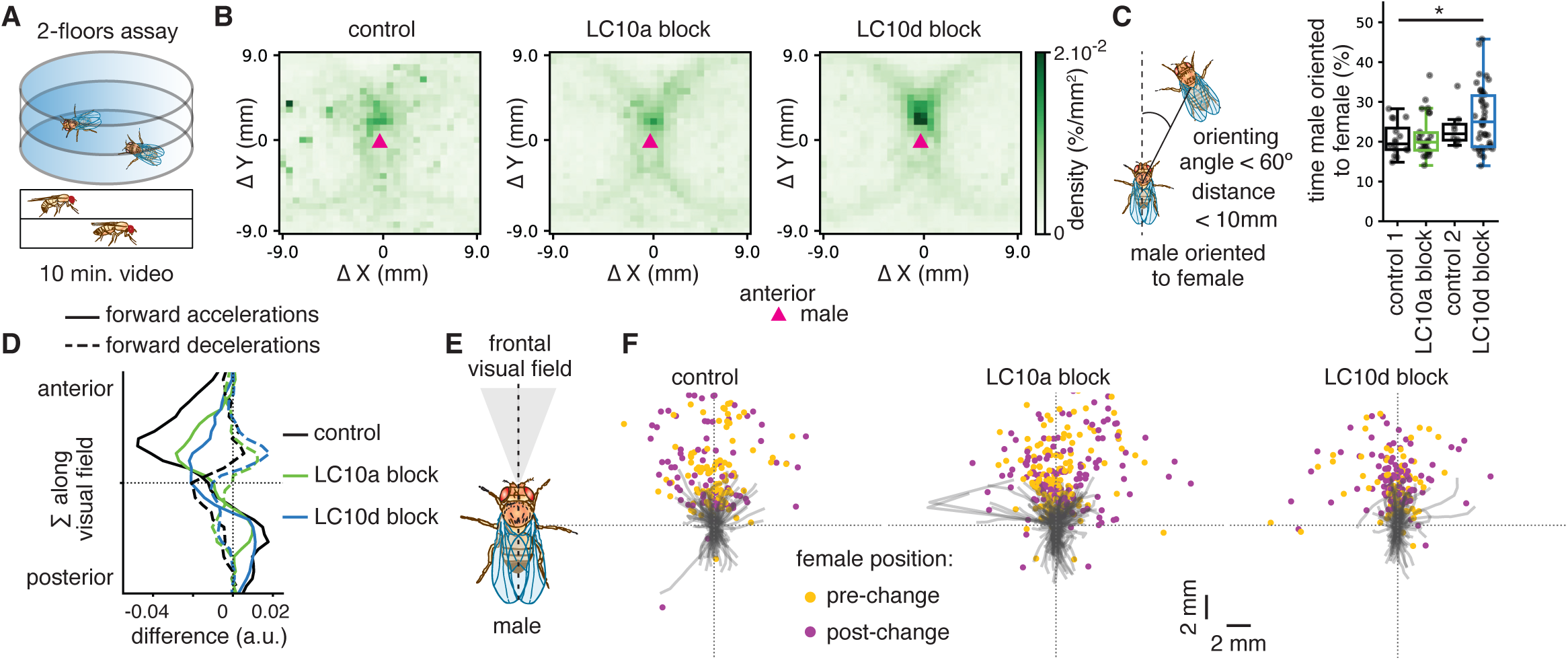
Avoidance in a vision-only assay. **A.** Schematic showing the chambers for the 2-floors assay based on Pan, et al, 2012. **B-F.** Parental control (N=16), LC10a-SS1 block (N=26), and LC10d-SS3822 block (N=35). **B.** Male centered heatmaps for female relative positions for all frames in 10-minute assays for controls, LC10a-SS1 block and LC10d-SS3822 block. **C.** Left: Schematic showing the male orienting to the female, with the limiting values for absolute orienting angle and distance indicated. Right: Percentage of time the male is oriented to the female. (*) p < 0.05, Student T-test. **D.** Projection along the Y axis of the heat map after – before, restricted to the frontal visual field (absolute orienting angle below 15° as shown in E), for forward velocity accelerations (solid lines) and decelerations (dashed lines). The dotted lines mark zero in both the X and Y axes. Controls (black), LC10a-SS1 block (green) and LC10d-SS3822 block (blue). Mean A.U.C. for the anterior visual field for accelerations with p = 0.012 for control vs LC10a block, p = 0.003 for control vs LC10d block; for decelerations with p > 0.05 for control vs LC10a, p = 0.031 for control vs LC10d block, Student T-test. **E.** Schematic showing the narrow frontal visual field, from -15° to 15° orienting angle. **F.** Male trajectories for a time envelope of 800ms centered around the change point (center, (0,0) for XY coordinates), for all forward velocity changes, with female relative positions at the start (yellow) and end (magenta) of the trajectory, for control, LC10a block and LC10d block.

Blocking neurotransmission in LC10d neurons in males led to an increase in the likelihood of female relative locations to the front of the male for the entire duration of the assay, that was absent from controls and LC10a block males, with LC10d-block males spending more time oriented towards the female (Figure 2B,C). Both forward accelerations and decelerations contributed to an increase in female frontal positions (Figure 2D). The observed alterations in avoidance behavior are independent from overall male locomotion capabilities (Figure S2O-Q). The presence of the female in a narrow frontal visual field, defined by absolute orienting angles below 15°, resulted in a shift to more lateral female positions in control and LC10a block males upon a change in male forward velocity (Figure 2E,F). Silencing LC10d neurons disrupted this behavior, leading to a higher chance of the female remaining in the narrow frontal visual field position after LC10d block males changed their forward velocity.

Collision avoidance is a visual behavior involving a reduction in walking speed of the observer fly as both the observer and the other fly (or visual object) move along intersecting paths (Tanaka and Clark, 2022; Zabala et al., 2012). To assess whether LC10d neurons contribute to collision avoidance, time points with inter-fly distances below 2 mm (collision points) were selected from the pre-courtship period of single pair assays or the entire duration of 2-floor assays (Figure S3A-I). Forward velocity for progressive or regressive object motion in the time leading up to the collision point displayed a mild deceleration upon object approach, assessed with increase in object angular size, in wild type, controls and LC10d block males (Figure S3C-E). A light reduction in forward velocity also preceded collision points in 2-floor assays for all genotypes (Figure S3F-I), suggesting that the mild collision avoidance present in our assays was unaltered by blocking LC10d neurons. Together, these observations implicate LC10d neurons in visual object avoidance that occurs towards objects devoid of a chemosensory profile, even if the visual object in question is a virgin female, and is independent of collision avoidance.

## Functional tuning properties of LC10d neurons

Among the six LC10-group neuron types identified to date, LC10a and LC10d are the most similar at the morphological level, with the main difference lying on the extent of the arborization to deeper layers in the lobula (Wu et al., 2016). The female and male connectomes show that there are 118 and 143 LC10a cells and 94 and 107 LC10d cells per hemisphere, respectively (Dorkenwald et al., 2024; Matsliah et al., 2024; Nern et al., 2025; Plaza et al., 2022; Scheffer et al., 2020; Schlegel et al., 2024; Seung, 2024; Zheng et al., 2018). The genetic driver LC10a-SS1 covers all LC10a cells, expressing on average in 150 cells per hemisphere, whereas the genetic access to LC10d neurons, LC10d-SS03822, targets roughly 75 cells per hemisphere (Figure S3J-N). Despite extensive similarities, LC10a and LC10d are two different neuron types based on genetic access, morphology and connectivity.

To determine the sensitivity of LC10d neurons, we used functional imaging of calcium transients detected with the calcium sensor GCaMP6m (Chen et al., 2013; Maisak et al., 2013), in regions of interest (ROIs) automatically segmented, and in response to a barrage of visual stimuli (Portugues et al., 2014; Ribeiro et al., 2018). All visual object stimuli, bright or dark, square- or long-bars, elicited a calcium response in LC10d lobula arbors, devoid of direction selectivity (Figure 3A-E).

**Figure 3.**
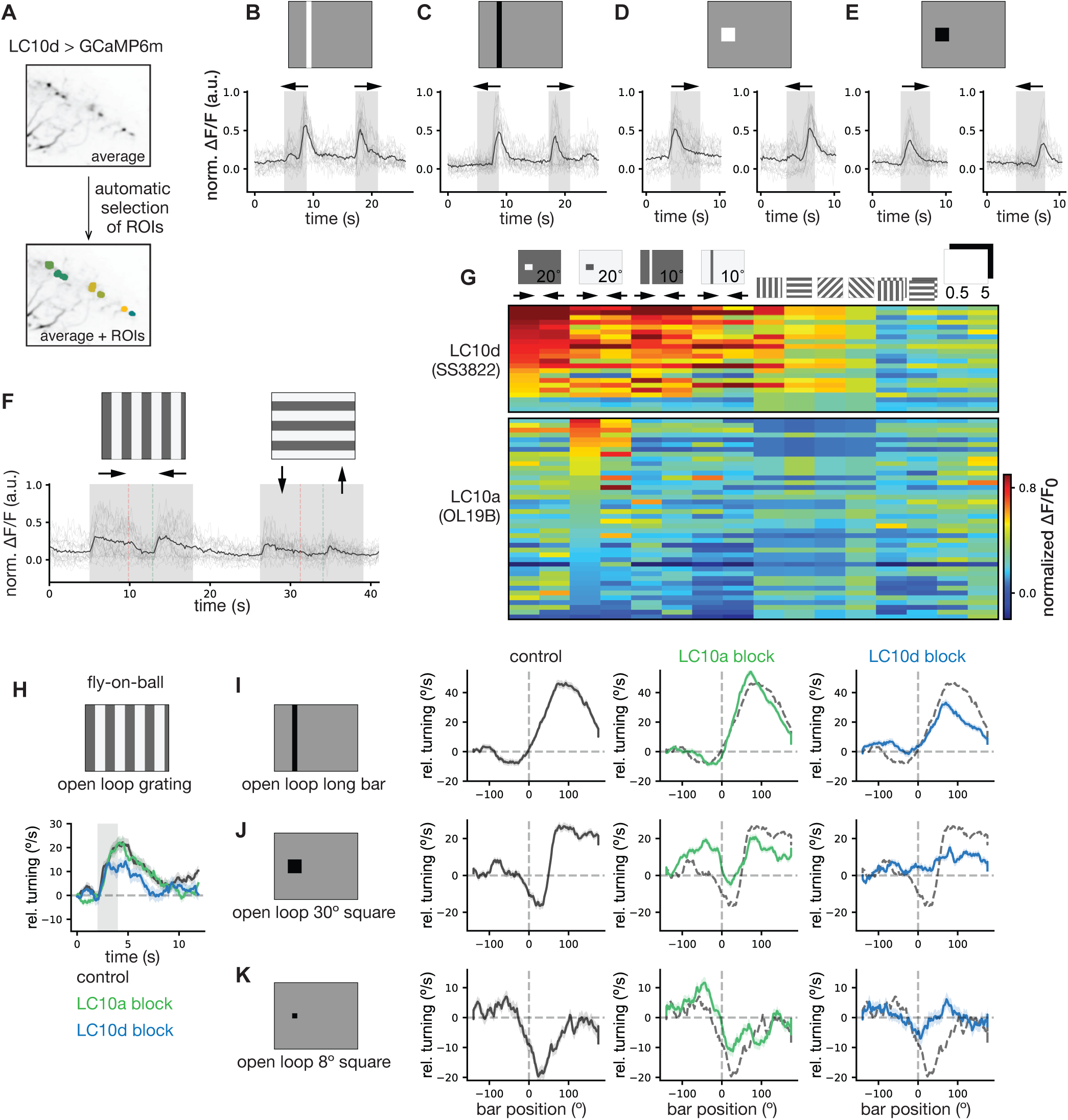
Tuning properties of LC10d neurons. **A.** Average image over time for one trial of presentation of all visual stimuli of LC10d-SS3822 lobula arborizations expressing GCaMP6m, showing automatic segmentation of regions of interest (ROIs). **B-F.** Calcium responses for *LC10d-SS3822 > GCaMP6m* (N=5), plotted as normalized change in fluorescence over base line fluorescence (ΔF/F_0_) for all ROIs (thin lines) and the mean (thicker line), in response to visual stimuli, shown in the shaded area of the time axis in the plot: long, 10° bar bright (B) and dark (C), square 30° bar bright (D) and dark (E), and gratings (F). G. Maximum ΔF/F_0_ per visual stimulus depicted on the top, for each ROI of LC10d-SS3822 (N=5 flies, same as in A-F) and LC10a-OL19B (N=3 flies). ROIs were sorted by the strongest response to bright square bar with clockwise direction for LC10d-SS3822 and to dark square bar with clockwise direction for LC10a-OL19B. **H-K.** Fly-on-ball assay to measure behavioral responses to visual stimuli for control (N=25, dark grey), LC10a-SS1 block (N=27, green) and LC10d-SS3822 block (N=25, blue). **H**. Relative turning over time for gratings. **I-K.** Relative turning in function of bar position for long, dark 10° bar (I), square, dark 30° bar (J) and small, dark 8° bar (K), for the indicated genotypes.

Presentation of bright and dark edges in eight different directions elicited calcium transients in all flies tested, without discernible direction selectivity (Figure S3O). Vertical gratings moving horizontally, and horizontal gratings moving vertically elicited low calcium responses in LC10d neurons (Figure 3F). Full field ON and OFF flicker pulsed every 5 seconds led to low amplitude calcium transients (Figure S3P), whereas negligible transients were observed in response to counter-phase flicker (Figure S3Q). Together, these observations indicate that LC10d neurons are broadly tuned to moving visual objects of different contrasts and sizes.

We compared LC10d tuning properties with those of LC10a neurons targeted with the OL19B driver and presented with the same visual stimuli (Figure 3G). LC10a-OL19B neurons exhibited strong responses to dark square-bars, devoid of consistent direction selectivity, followed by responses to bright square-bars and full field flicker, similar to what was previously reported (Ribeiro et al., 2018; Schretter et al., 2025). In contrast, LC10d-SS3822 neurons displayed larger amplitude calcium transients in general and were sensitive to long bars and gratings in addition to square-bars and full field flicker. LC10d neurons are thus more broadly tuned than LC10a neurons. These observations show that the AOTU receives rich and diverse visual information, as predicted by the connectome (Matsliah et al., 2024; Nern et al., 2025; Seung, 2024), that includes at least two object detecting neurons, LC10a and LC10d.

Upon entry into a persistent courtship or aggression internal states, LC10a neuronal activity is increased as a result of dendritic disinhibition, and a toggle switch motif (Hindmarsh Sten et al., 2021; Rubin et al., 2025; Schretter et al., 2025). We reasoned such alterations reported in LC10a responses might be detected with a reporter that integrates neuronal activity occurring on the scale of minutes, like CRTC (Bonheur et al., 2023). We leveraged the 2-floor arenas to compare LC10a activity in the 2-floor, vision-only setting, in which the male is exclusively exposed to female visual cues (Figure S4A,B), with LC10a activity during courtship, by placing the male and female in the same floor and in which the male is exposed to all sensory cues from the female (Figure S4C,D). Female tracking is absent in vision-only conditions (Figure S4B), but extensive when the male is exposed to all female sensory cues and courtship occurs (Figure S4D). The distribution of CRTC signal was skewed to the nucleus in LC10a neurons in males that were exposed to the entire female sensory cocktail, but not in males exposed solely to visual cues, as observed by the overlap of CRTC:EGFP with the nucleus marker DAPI compared to CRTC:EGFP overlap to the plasma membrane marker mCherry:CD8 (Figure S4E,F). In contrast, CRTC:EGFP distribution to the nucleus and cytoplasm of LC10d neurons remained the same with exposure to either the entire cocktail or solely visual sensory cues from the female (Figure S4E,F). The LC10d neuronal activity thus remained unchanged upon courtship initiation, suggesting that LC10d neurons escape modulation by the persistent courtship internal state.

Tethered males walking on an air-suspended Styrofoam ball were shown visual stimuli on three computer screens surrounding the fly (Bahl et al., 2013). The air keeping the ball afloat was kept at 25°C, the preferred temperature of *Drosophila melanogaster* (Sayeed and Benzer, 1996). Male locomotion was assessed with relative turning, in which positive values indicate turning in the same direction as the stimulus object, whereas negative values indicate turning in the opposite direction of the stimulus object. Males with LC10a or LC10d neurons blocked displayed optomotor response to full field gratings, and tracked a long, dark bar similar to control males (Figure 3H,I), albeit with lower amplitudes for relative turning in the case of LC10d block males. Responses of control males to a 30° dark, square bar involved reduced turning during back-to-front relative motion, a brief turning away when the bar traversed the frontal visual field with front-to-back relative motion, and turning towards the bar during front-to-back motion in the last stretch of the trajectory in the visual display across more lateral positions (Figure 3J). This profile of turning was altered in LC10a block males, with turning towards the bar occurring with back-to-front relative motion, and absence of turning away when the bar crosses the frontal visual. This suggests that LC10a block males detect the presence of the 30° bar but are unable to track it. Blocking LC10d neurons led to reduced turning amplitude for the entire bar trajectory, with the direction of turning relatively similar to that of controls. Presentation of an 8° dark, square bar led to a relative turning against the bar movement in both control and LC10a block males (Figure 3K). Turning away from the 8° bar was severely reduced in LC10d block males, suggesting that avoiding of a small bar relies on LC10d neurons. Importantly, bar angular sizes remained constant, suggesting that avoidance of small visual objects occurs in the absence of looming and differs from collision avoidance (Tanaka and Clark, 2022; Zabala et al., 2012). We conclude that LC10d neurons mediate visual object avoidance in tethered walking in the fly-on-ball paradigm, and contribute to tracking long and square bars to a lesser extent.

### LC10a and LC10d coordinate diverse motor outputs

The LC10a and LC10d terminals share several common downstream AOTU-output neuron types (Figure 4A,B), according to the female and male whole brain connectomes (Berg et al., 2025; Dorkenwald et al., 2024; Plaza et al., 2022; Schlegel et al., 2024; Zheng et al., 2018). Paradoxically however, we found that LC10a and LC10d neurons are required for opposing behaviors in response to visual objects, tracking and avoidance respectively. Based on these findings, we posit that unilateral, optogenetic activation of LC10a or LC10d might elicit diverging motor outputs. The LOV-LexA light-gated gene expression tool (Figure 4C) was used to express the cation channel CsChrimson in groups of two to seven cells of LC10a or LC10d neurons in one brain hemisphere (Klapoetke et al., 2014; Ribeiro et al., 2022), and behavior in single flies was assessed. Similar to previous reports (Hindmarsh Sten et al., 2021; Ribeiro et al., 2018), activation of unilateral LC10a cells resulted in turning to the same side of expression (Figure S4G-I). The cumulative relative rotational velocity was clearly skewed to the ipsilateral side, even if the number of cells unilaterally activated remained under seven (Figure 4D). In contrast, unilateral activation of LC10d neurons led to ambiguous modulation of rotational speed (Figure S4J-L), that was unbiased towards ipsi- or contralateral side of expression (Figure 4E). Unilateral activation of both LC10a and LC10d neurons elicited a modest increase in forward velocity (Figure S4I,L), that was absent in flies bearing no expression of CsChrimson (not shown). These observations suggest that, despite sharing several common downstream neuron types, LC10a and LC10d impinge different motor outputs on fly locomotion, likely through different downstream pathways.

**Figure 4.**
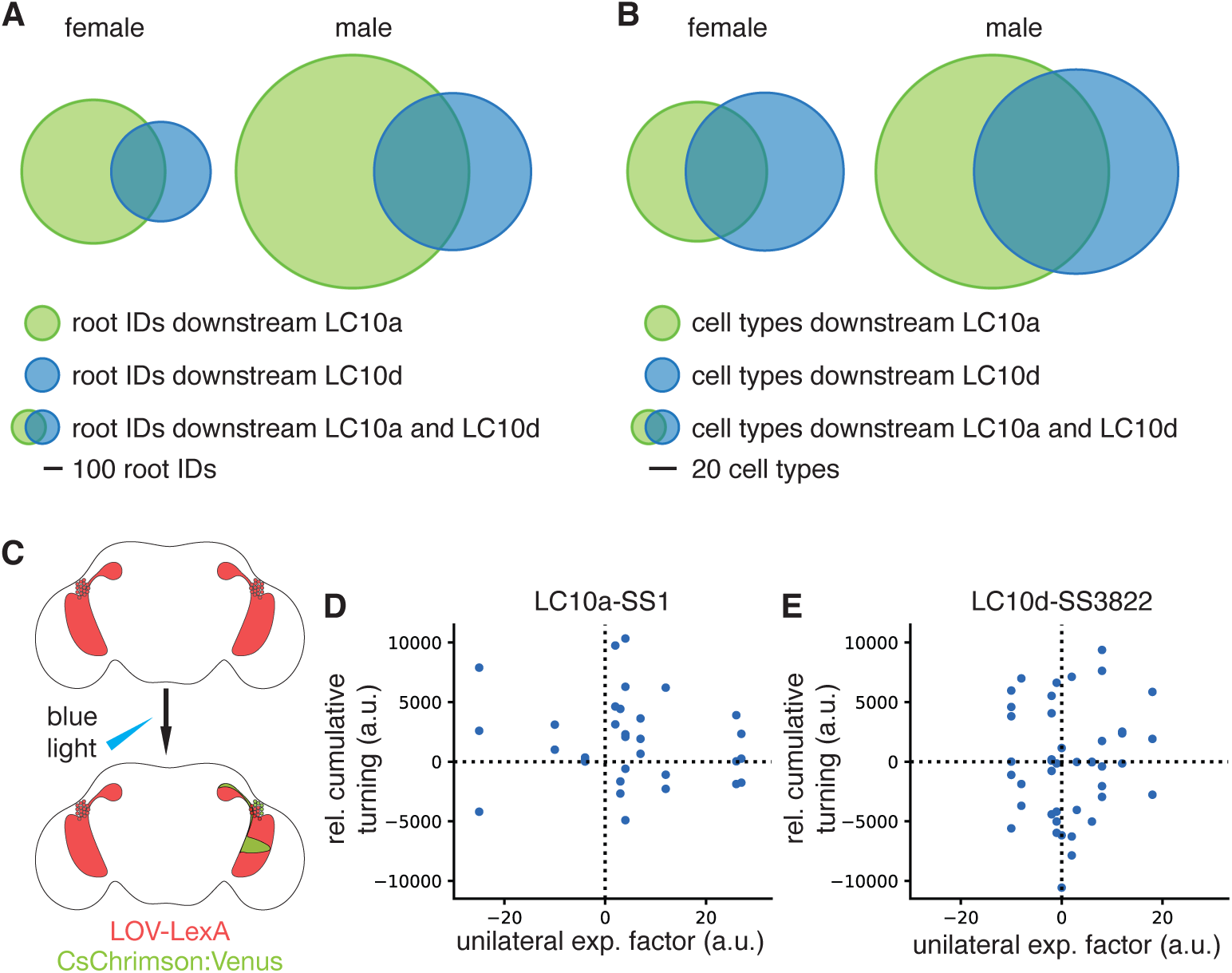
Circuit and behavioral outputs of LC10a and LC10d. **A.** Plot showing the proportion of neuronal cells (root IDs) downstream all LC10a and LC10d neurons, in the female (left) and male (right) connectomes. All circle diameters are proportional to the number of downstream root IDs for LC10a neurons in the male connectome. **B.** Plot showing the proportion of consolidated neuron types downstream all LC10a and LC10d neurons, in the female (left) and male (right) connectomes. All circle diameters are proportional to the number of the downstream unique neuron types for LC10a neurons in the male connectome. **C.** Schematic showing light-gated expression of CsChrimson:Venus with LOV-LexA. **D, E.** Cumulative relative turning in function of unilateral expression for all flies tested for LC10a- SS1 > LexA-LOV, CsChrimsonVenus (N=12, D), and LC10d-SS3822 > LexA-LOV, CsChrimsonVenus (N=16, E).

### Chunking of the visual field underlies LC10d-dependent object avoidance

A handful of neuron types receive more input from LC10d than from LC10a neurons (Figure 4A,B). Among them is the descending neuron DNa10, composed of a single cell per hemisphere, that is downstream of roughly one third of ipsilateral LC10d cells both in males and females (Figure 5A,B). In addition to LC10d, DNa10 neurons receive inputs from LLPC3 neurons and to a lesser extent, from LPLC4 and other LC10-group neurons (Figure S4P). Remarkably, the LC10d subpopulation upstream DNa10 resides in a region in the lobula that represents the anterior or frontal visual field (Figure 5C, S4M,N) (Dorkenwald et al., 2024; Schlegel et al., 2024; Zhao et al., 2025; Zheng et al., 2018). A handful of neuron types also receive inputs preferentially or exclusively from the anterior facing LC10d subpopulation, albeit at lower levels compared to DNa10 (Figure 5D). LC10d-visual information is thus organized in downstream circuits into zones with likely different ethological relevance.

**Figure 5.**
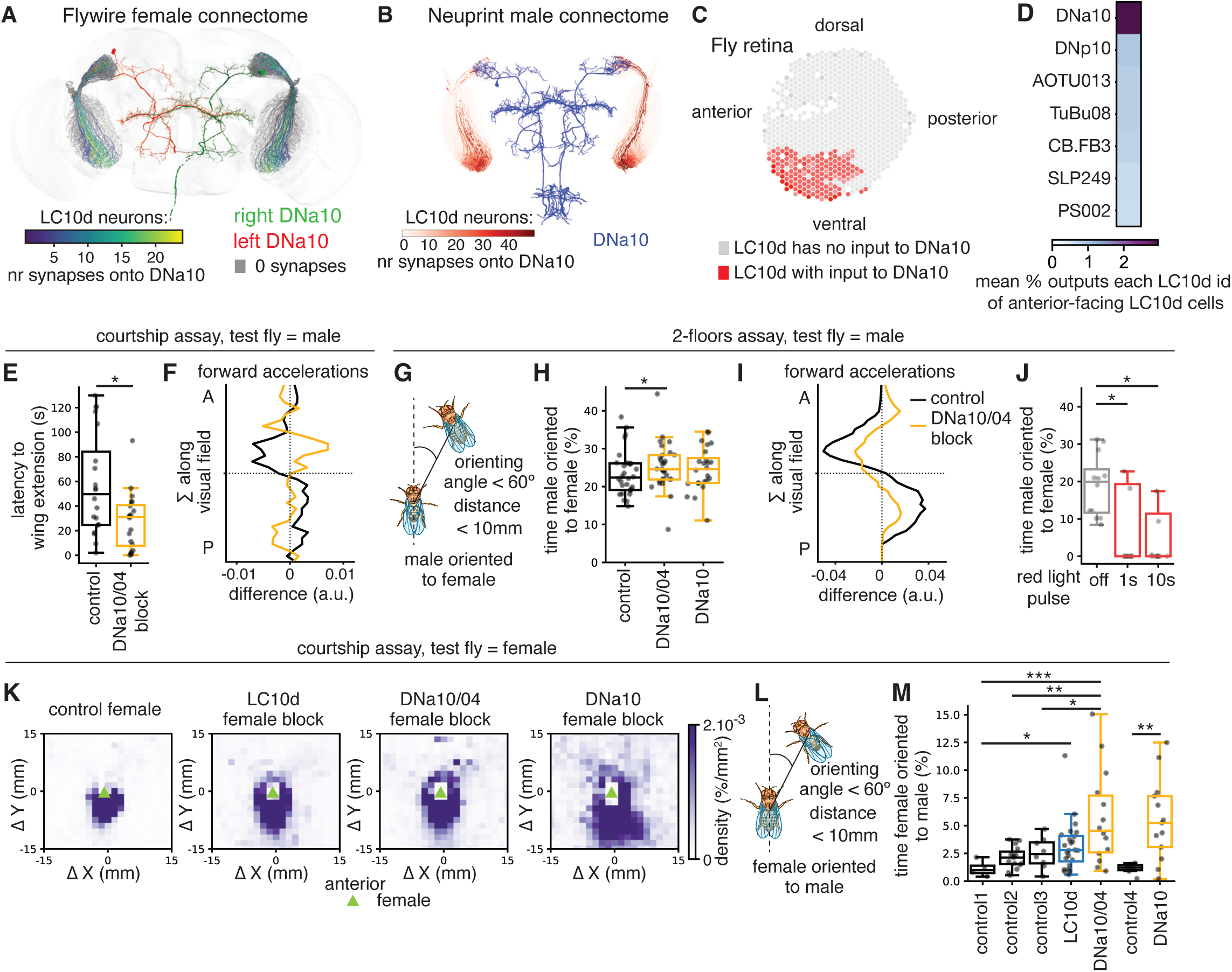
Chunking of spatial information underlies LC10d-dependent avoidance. **A, B.** LC10d – DNa10 connection based on the female (A, FAFB) and the male (B, MCNS) connectomes. **C.** Planar projection of ommatidia in the retina, with ommatidia upstream LC10d cells that synapse onto DNa10 highlighted in red. **D.** Mean percentage of outputs per LC10d cell for downstream neuron types that receive more synapses from anterior facing LC10d neurons, based on the male connectome MCNS. **E-F.** Single pair courtship assay in 42mm arenas in which the male is the test fly, for parental control (N=20) and DNa10/04-SS2384 block (N=20). **E.** Latency to initiate courtship as determined by the first single wing extension, p < 0.05 for control vs DNa10/04-SS2384 block, Student T-test. **F.** Projection of the female relative positions along the male visual anterior-posterior axis resulting from the difference after-before for male forward accelerations. Mean A.U.C. for the anterior visual field with p > 0.05 for control vs DNa10/04-SS2384 block, Student T-test. **G-I.** Vision-only 2-floors assay in which the male is the test fly, for parental control (N=30), DNa10/04-SS2384 block (N=29), and DNa10-SS70607 (N=24). **G-H.** Schematic of the definition of male oriented to the female (G) and percentage of time the male is oriented to the female (H). (*) p < 0.05, Student T test with outliers removed. **I.** Projection of the female relative positions along the male visual anterior (A)-posterior (P) axis resulting from the difference after-before for male forward accelerations. Mean A.U.C. for the anterior visual field with p = 0.02 for control vs DNa10/04-SS2384 block, Student T-test. **J.** Optogenetic activation of DNa10 neurons in males (DNa10/04-SS2384 > UAS- CsChrimson:Venus, N=12) paired with wild type females in 2floor assays, with three trials for each light pulse per pair. (*) p < 0.05, Student T-test. **K-N.** Single pair assay, in which the female is the test fly, for parental control (N=15); LC10d-SS3822 block (N=14); DNa10/04-SS2384 block (N=14); and DNa10-SS70607 (N=13). **K.** Female centered heat maps of male relative positions for the courtship assay. **L.** Schematic showing the definition of female oriented to the male. **M.** Percentage of time the female is oriented to the male. (*) p < 0.05, (**) p < 0.01, (***) p < 0.001, Student T-test.

Since DNa10 receives significantly more input from LC10d than LC10a neurons, in males and females (Figure S4O), we hypothesized that DNa10 might mediate avoidance of visual objects when these are located in the frontal visual field of the observer fly. Two driver lines were used to access DNa10 neurons, SS02384 which also targets DNa04 (DNa10/04-SS2384) and SS70607 (DNa10-SS70607) (Meissner et al., 2024; Namiki et al., 2018). Males with DNa10/04-SS2384 blocked initiated courtship significantly earlier in single pair assays (Figure 5F), while blocking DNa10-SS70607 neurons led to a tendency to early courtship initiation that was not significant (not shown). Similar to LC10d block males, accelerations in male forward speed led to a mild increase in the likelihood of anterior female relative positions upon DNa10/04-SS2384 silencing (Figure 5F). In the 2-floor assay, silencing DNa10/04-SS2384 or DNa10-SS70607 neurons in males resulted in an increase in the percentage of time the male is oriented to the female compared to controls (Figure 5G,H). In addition, male forward accelerations led to a shift in female relative positions to the frontal visual field, that is significantly different from controls (Figure 5I). A small decrease in walking speed was observed in the time leading to collision in controls, DNa10/04-SS2384 and DNa10-SS70607 block males in both assays (Figure S5A-L and not shown). Optogenetic activation of DNa10/04-SS2384 neurons resulted in a decrease in the percentage of time the male is oriented to the female in 2-floor assays (Figure 5J). These observations imply that the anterior facing LC10d cells mediate avoidance through the DNa10 descending neuron in males.

When initially placed in the courtship arena, both the male and the female appear to explore the new space, displaying little interest towards the other fly. Upon arousal to court, male behavior switches to orienting and chasing the female while singing. Changes in female behavior are more elusive. Her continuous locomotion may indicate that the female simply continues to explore the arena. However, the observed reduction in female forward speed after courtship starts, suggests that female locomotion is also actively regulated (Coen et al., 2014). Male positions relative to female-centered coordinates were highly enriched in the posterior field of the female for the entirety of the single-pair assay (Figure 5K) (Agrawal et al., 2014). Silencing LC10d or DNa10 neurons in females led to an enrichment of male positions frontolateral relative to the female (Figure 5K), with LC10d, DNa10/04-SS2384 or DNa10-SS70607 block females spending more time oriented towards the male (Figure 5L,M). Interestingly, these alterations in female behavior led to a delay in initiation of male courtship behavior, with the latency to the first wing extension and following towards females with LC10d, DNa10/04-SS2384 or DNa10-SS70607 neurons blocked increasing significantly (Figure S5M,N and not shown).

The profile of female locomotion changes leading to frontolateral male positions relative to the female, displayed a similar overall trend to that of non-courting males: female forward accelerations and decelerations biased male relative positions towards the anterior visual field (Figure S5O), that was largely independent of collision avoidance in control, LC10d, and DNa10-SS70607 block females (Figure S5G-J,L). Blocking DNa10/04-SS2384 neurons in females resulted in elevated walking speeds that were stable in the time leading up to collision points (Figure S5K). The amplitude of forward and rotational velocities at change points, as well as their temporal profile, were similar between controls, LC10d-and DNa10/04-SS2384-block females (Figure S5P-R). Female locomotion during courtship is thus purposeful in reducing the likelihood of male relative positions from the female anterior visual field. Together, these observations indicate that the anterior looking LC10d sub-population of cells act via DNa10 to regulate locomotion maneuvers that reduce the likelihood of a visual object remaining in the frontal visual field of the observer fly.

## Discussion

It is advantageous to keep unknown objects at a safe distance to avoid harm in case the unknown object turns out to be nefarious, like a predator or a competitor. Our work uncovered LC10d neurons as mediators of avoidance of discrete objects that are perceived visually without a chemosensory signature or tell-tale cues of an imminent attack. LC10d neurons arbitrate locomotion maneuvers that reduce the likelihood of frontal positions of the visual object relative to the observer fly in several contexts: during a roughly one-minute long pre-courtship period that occurs in large arenas in single-pair assays, and precedes integration of fly chemosensory cues; during male – male pair assays in large arenas in the absence of food; during exposure to naturalistic visual cues alone from the female in the vision-only, 2-floor assay; during exposure to small bars in the fly-on-ball set up; and in female behavior during courtship. LC10a and LC10d are object detecting neurons that mediate opposing behaviors and display different sensitivity to the courtship internal state, implying that LC10a and LC10d downstream circuits function in different sensory contexts to provide increased flexibility in behavioral responses to objects. We further show that DNa10 descending neurons, downstream of a LC10d sub-population representing the anterior visual field, mediate avoidance in the same behavioral paradigms, through regulation of forward accelerations and decelerations that is similar to LC10d neurons.

A defining characteristic of the AOTU is the spatial organization of its input LC neuron types (Hoeller et al., 2025; Nern et al., 2025; Plaza et al., 2022). Both LC10a and LC10d neurons are composed of dozens of cells per hemisphere, that display coarse retinotopy in their terminals in the central unit of the AOTU (Ribeiro et al., 2018; Wu et al., 2016). In contrast, AOTU central unit output neuron types are typically composed of far fewer numbers of cells, that are poised to synthesize information from subsets of cells of LC10-group neuron types (Berg et al., 2025; Dorkenwald et al., 2024; Plaza et al., 2022; Schlegel et al., 2024; Zheng et al., 2018). Our work identifies chunking of visual field space as a mechanism of synthesizing AOTU-central unit visual inputs. A single cell of the descending neuron DNa10 receives inputs from just under one third of LC10d cells of the same hemisphere, that collectively represent the anterior visual field(Berg et al., 2025; Dorkenwald et al., 2024; Keleş and Frye, 2017; Namiki et al., 2018; Scheffer et al., 2020; Schlegel et al., 2024; Zheng et al., 2018). This one third, anterior field LC10d sub-population appears to have a starker influence on avoidance behavior in the paradigms tested here, since blocking DNa10 neurons produces similar phenotypes in males and females to blocking most LC10d cells. In addition, activation of DNa10 neurons elicits avoidance. Our work thus provides evidence that the LC10d – DNa10 circuit chunks the anterior visual field of the observer fly to regulate object avoidance.

In the ventral nerve cord, DNa10 neurons mainly project to the wing, intermediate and lower tectulum regions, involved in wing control and coordination between wing and leg movements (Cheong et al., 2026). A smaller proportion of DNa10 output synapses locate to leg neuropils. Although social interactions in *Drosophila melanogaster* occur mostly during walking on a substrate, wings are involved in courtship song production and several wing movements in aggression (Coen and Murthy, 2016; Vrontou et al., 2006), in addition to escape and flight. DNa10 neurons might be implicated in regulation of two different behaviors: steering maneuvers in specific contexts as shown here, and control of wing motion for social communication or flight. Differences in activation thresholds in neurons downstream of DNa10 could underlie such dual roles. The descending neurons DNa02 control turning during walking (Rayshubskiy et al., 2025), and although most DNa02 projections go to leg circuits, a smaller proportion indirectly connects to one wing motoneuron in the ventral nerve cord (Cheong et al., 2026). Further research is necessary to understand the roles of concomitant projections of DNs to wing and leg circuits.

DNa10 receives visual information from LC10d neurons representing the anterior visual field. A similar organization of visual space occurs in two LC10a downstream neurons, AOTU019 and AOTU025 (Collie et al., 2026). The lateral zone of the visual field is more represented in LC10a-inputs to AOTU025, whereas the anterior or frontal zone is more represented in LC10a-inputs to AOTU019. Accordingly, AOTU025 neurons are active during orienting maneuvers towards an object stimulus, whereas AOTU019 neurons are required to maintain the object in the narrow frontal visual field of the pursuer. Moreover, division of the visual space into zones is common in AOTU-output neurons (Collie et al., 2026; Hoeller et al., 2025), suggesting that chunking the observer’s field of view into zones with different ethological meaning is a fundamental organizational principle of AOTU-downstream pathways.

The life ecology of Drosophila species involves co-habitation of fruits with many flies of the same species as well as other Drosophila species (Markow, 2015), naturally leading to the simultaneous detection of many discrete objects of variable sizes, with various chemical signatures. Several visual projection neuron types are likely to guide the observer fly in moving in such environments. At least sixteen LC neurons are sensitive to discrete objects in Drosophila (Aptekar et al., 2012; Keleş and Frye, 2017; Klapoetke et al., 2022; Ribeiro et al., 2018; Schretter et al., 2025; Städele et al., 2020). In addition to a considerable overlap in their tuning properties, as assessed with current stimuli, visual information from the same object detecting neuron can be used to dissect diverse visual scenes. LC11 neurons mediate collective detection of several flies in an appraisal of safety cues to buffer defensive behavior and, in the presence of a single small visual object, contribute to halting (Ferreira and Moita, 2020; Tanaka and Clark, 2020). In contrast, LC10a and LC10d neurons display similar sensitivities to visual objects, yet participate in different behaviors. We show that LC10d activity levels are indifferent to the courtship internal state, contrary to LC10a, whose activity is enhanced upon courtship initiation (Hindmarsh Sten et al., 2021; Ribeiro et al., 2018; Rubin et al., 2025; Schretter et al., 2025). In this case, top-down control is used to disentangle similar visual information in the central brain. The elevated number of object detecting neurons in the fly brain and their seemingly overlapping sensitivities suggest that more circuit mechanisms are bound to play a role in portraying the visual environment in Drosophila in their natural habitat, including chunking of the visual space.

Visual information conferred by mammalian retinal ganglion cells is further processed in the superior colliculus and in other deeper brain regions that are linked to several behaviors (Allen et al., 2021; Isa et al., 2021; Monavarfeshani et al., 2017). In zebrafish, prey pursuit and social affiliation are associated with distinct areas of the optic tectum, that encode visual information imbued with ethological significance (Helmbrecht et al., 2018; Kappel et al., 2022; Larsch and Baier, 2018; Semmelhack et al., 2014). In insects, visual information is often ethologically relevant at the level of visual projection neurons since optogenetic activation of several LC neuron types can elicit specific behaviors (Wu et al., 2016). We show that LC10d neurons relay broad visual object information to the central brain apparently detached from any valence, since unilateral activation of LC10d neurons did not bias direction of locomotion. LC10a neurons are targets of top-down modulation, and encode visual objects deemed to be attractive by a persistent internal state dependent modulation, with unilateral activation leading to ipsilateral turns (Bath et al., 2014; Hindmarsh Sten et al., 2021; Inagaki et al., 2014; Ribeiro et al., 2018; Schretter et al., 2025). The function of LC10d neurons might be to simply encode objects present in a visual scene, with the choice of the appropriate behavioral response to those objects lying on other visual projection neurons or downstream neural circuits.

The AOTU central unit receives diverse visual information from six different LC neuron types. Despite sharing several common downstream neurons, we show that LC10a and LC10d participate in at least two different behaviors, tracking and avoidance. This suggests the AOTU central unit functions as a hub for processing visual information that contributes to diverse behaviors in Drosophila. The LC10d sub-population representing the anterior visual field and the downstream DNa10 neurons chunk visual space into a zone that is relevant for avoiding a potentially harmful interaction, showcasing one possible synthesis of the map-like spatial information represented in the central unit of the AOTU.

### Limitations of this study

Single-pair assays are an incomplete emulation of the entire repertoire of interactions among Drosophila flies living on a food substrate. The high throughput of such behavioral paradigms however enables the study of many more neuron types and conditions on the already immensely complex social interactions between two flies. Our level of behavioral analysis offers a glimpse into detection of a single other fly or another animal in the distance. The neural circuit here uncovered might participate in more behaviors occurring in the presence of several other flies. Testing behavior in smaller arenas also increases throughput, but exposes flies to limited sets of visual cues. The consequences to a behavior occurring in close proximity, like courtship, are negligible. However, avoidance of distant animals might be affected in confined spaces. Lastly, we used TNT as the single effector to block neurotransmitter release, since TNT has been corroborated by many studies. We cannot exclude however, that chronic silencing of neurotransmission might be more relevant in our assays than acute manipulations with optogenetic tools.

## Materials and Methods

**Table S1**

List of genotypes per figure.

**Table S2**

List of root cell IDs used per neuron type.

### Contact for Reagent and Resource Sharing

Please contact Inês M.A. Ribeiro, at.

### Experimental Model and Subject Details

*Drosophila melanogaster* strains used in this work are listed in the Table S1. Flies were raised in vials containing standard cornmeal-agar medium supplemented with baker’s yeast and incubated at 25°C with 60% humidity and 12h bright light/dark cycle throughout development and adulthood, unless otherwise noted. Both males and females were the subjects of interest in this work. For single pair assays in 18- or 42-mm arenas, and 18-mm 2-floor assays, males were collected two to three hours after emergence from the pupal case and single housed in a 96-deep well plate with 0.5 ml of unsupplemented standard cornmeal-agar medium. This treatment keeps males socially naïve. The females were collected as virgins, also two to three hours after pupal emergence, and group housed with other virgin females. Males and females were collected as non-virgin and group housed together for functional imaging and optogenetic activation experiments. Both males and females used in behavioral experiments, functional imaging or dissections between the ages of 4 to 9 days old. Neurotransmission was blocked in specific neurons with tetanus toxin light chain (TNT) (Sweeney et al., 1995).

### Behavioral assays

Single-pair assays were conducted within three hours after the incubator lights turned on, in ZT 0 to 3 hours. A plate held two 42-mm barrierless arenas, but video recording was done one at a time. One naïve male and one virgin female were gently sucked separately into the arena, the plate placed roughly 20 cm underneath a camera and illuminated with abundant bright light from the sides. For figure 1, a JVC HD video camera was used, with a resolution of 1440x1080 (16:9) at 25 fps for 10 min. For assays in 42-mm arenas in figure 5, S1K-L, and S5, videos were recorded with FLiR cameras with a resolution of 1280x1024 at 60 fps for 10 min or until copulation started. Videos were acquired in the .MTS or .AVI file format, without rendering, tracked with the custom software MateBook version 1241 (Ribeiro et al., 2018). XY coordinates of fly centroid and head, classifiers as well as locomotion parameters were used for data analysis.

For the 2-floors assay, two plates, each holding eight arenas with a diameter of 18 mm and height of 3 mm, were stacked on top of each other and separated by a transparent acrylic plate with 1 mm thickness. Video recording was initiated immediately after all the flies were loaded into the 2- floor chambers, which typically lasted less than one minute. Filming was performed with a Raspberry Pi Global Shutter Camera and a Raspberry computer with a resolution of 1456x1088 and 30 fps. The video format .H264 was reformatted into .MP4 and tracked with MateBook version 1241. Further analysis of behavioral data was carried with Python 3.2 with the libraries NumPy, SciPy and Matplotlib. Scripts are available upon request. All behavioral assays were conducted between 24 to 28°C, with 50% humidity.

### Unbiased analysis of fly locomotion

The software MateBook provides time series for X and Y coordinates of center of the ellipse fitted to the male or the female (fly centroid, located roughly at the intersection of the thorax and abdomen), from which forward and rotational velocities are extracted. To find the timepoints at which the male changed his locomotion speed, the derivative of the forward velocity was scanned every 10 frames (for 25 fps videos) or 400 ms for local maxima and local minima, using as cutoff two times the standard deviation of the derivative. Each local peak was attached with 400 ms before and after, and this trace scanned for presence of missegmented frames. Traces were kept for further analysis if the number of missegmented frames did not exceed 15 for the entire trace (in 25 fps videos). Local maxima traces represent accelerations and local minima represent decelerations. The same analysis was completed for rotational velocity. However, elevated turning speeds in our behavior paradigms had very short durations and local maxima were invariably followed by local minima within 40 ms. We thus considered all local peaks for rotational velocity together, without differentiating between accelerations and decelerations.

### Assessment of neuronal activity

Functional imaging with a custom built 2-photon confocal microscope was performed as previously (Maisak et al., 2013; Ribeiro et al., 2018). Briefly, males or females were glued to a recording chamber, immobilized with bee wax, leaving the right eye free. The recording chamber was filled with Ringer solution (103mM NaCl, 3mM KCl, 5mM TES, 10mM trehalose, 10mM glucose, 3-7mM sucrose, 26mM NaHCO3, 1mM NaH2PO4, 1.5 CaCl2 and 4mM MgCl2, pH 7.3 to 7.35, 280-290mOsmol/Kg) (Mauss et al., 2014). A surgery was performed to open a window in the back of the right eye and to expose the lobula. The calcium sensor GCaMP6m (Chen et al., 2013) was expressed in LC10d with the SS3822 driver and in LC10a with the OL19B driver (Wu et al., 2016). All recordings were made in a w+ background. The visual stimuli were presented in a projector based arena and were previously described (Arenz et al., 2017; Ribeiro et al., 2018).

The CRTC:EGFP (Bonheur et al., 2023) tool was used to assess integration of minute long neuronal activity in two conditions: in the vision-only, 2-floors assay, in which the male and female are placed in two different, stacked arenas that are separated by a transparent acrylic plate (Figure S4A); in the courtship assay, in which the male and female are placed in the same arena (Figure S4C). The behavior was video recorded for 10 minutes, after which the flies were cold anesthetized, dissected and immunostained as previously described (Ribeiro et al., 2018), with the primary antibodies: anti-GFP (chicken, 1:1000 dilution), anti-RFP (rabbit, 1:1000 dilution), and the secondary antibodies anti-chicken Alexa488 and anti-rabbit Alexa568. Finally, the brains were incubated in 1:3000 DAPI in PBST for five minutes, washed and mounted on a slide in Vectashield. The brains were imaged with a Leica SP8 with a 20x oil immersion objective (Leica Immersion 20x/0.7).

### Optogenetics

Males aged between 5 and 7 days, expressing the cation channel CsChrimson:Venus with DNa10-SS2384 driver and fed *all-trans retinal* throughout adulthood (30mg/ml in yeast paste), were tested in the top arena of 2-floor assay arenas and paired with a wild type female in the bottom floor. The 2- floor assay fly pairs were video recorded in dim blue light, and provided with red light at 3.8 mW/cm^2^ for 1 or 10 seconds. The videos were tracked with MateBook and the behavioral data was further analyzed with NumPy and MatPlotLib in Python 3.2.

By combining LexAop-CsChrimson:Venus (Klapoetke et al., 2014) and LOV-LexA (Ribeiro et al., 2022) it was possible to limit expression of CsChrimson:Venus up to seven LC10a or LC10d cells in one brain hemisphere. Light delivery to gate expression with LOV-LexA and optogenetic light exposure in the behavioral assay were performed as previously reported (Ribeiro et al., 2022, 2018).

Males and females between 4 and 6 days old, expressing LOV-LexA in LC10a-SS1 or LC10d-SS3822, reared at 18°C with limited light exposure, and provided with *all-trans retinal* throughout adulthood (30mg/ml dissolved in yeast paste), were anesthetized with CO2 and glued onto a plastic plate with four 300 to 400 µm diameter holes with Eicosene 99% (Aldrich 219274-5G). Each of four flies was fitted under one of the holes such that roughly one third of the head was centered on the hole. The rest of the fly body and head were shielded from light. The plastic plate was then mounted onto a glass slide with plasticine for height to accommodate the flies. Blue light delivery was performed with an upright confocal (Leica TCS SP8) with a preprogrammed serial delivery with the 458-nm laser at 2.49 µW, 0.33 Hz for 90 seconds, four times with an interval of 20 minutes. The flies were then detached from the plastic plate and the leftover Eicosene was removed from the fly thorax. Behavioral assays in 18-mm beveled chambers were performed roughly 24 hours later, following a protocol previously described (Ribeiro et al., 2018). Briefly, flies were filmed in dim blue LED light with a JVC camera and delivered red light for 3.5 seconds at 3.8 mW/cm^2^, for three trials. The .MTS videos were tracked with MateBook and SLEAP, which gave the same results. The behavior data in figures 4 and S4 is from MateBook and was plotted using Python 3.2 packages.

After filming the behavior, each fly was dissected, the brain incubated for 5 minutes with DAPi, mounted and imaged with a confocal (Ribeiro et al., 2022). The expression pattern in the brain of CsChrimson:Venus was analyzed with respect to the number of cells in either hemisphere and expression levels. The unilateral expression factor (unilateral exp. factor in figure 4D,E) was calculated based on the number of cells expressing CsChrimson:Venus on both hemispheres, multiplied by a factor representing expression strength, measured manually based on brightness of native fluorescence of CsChrimson:Venus in confocal images. The hemisphere with the highest number of cells was called ‘ipsilateral’, and the cumulative turning awarded a positive value if towards the ipsilateral side (the same side of the highest expression), or a negative value if towards the contralateral side. In short:

> unilateral exp. factor = (nr cells ipsilateral – nr cells contralateral) * expression strength

### Immunostaninig

Immunostaining for Venus and tdTomato in figure S3 was performed as previously described (Ribeiro et al., 2018; Yu et al., 2010). Briefly, anti-GFP (chicken, 1:1000 dilution) and anti-RFP (rabbit, 1:1000 dilution) antibodies were used to detect Venus and tdTomato respectively, along with the secondary antibodies anti-chicken Alexa488 and anti-rabbit Alexa568. The anti-TNT (goat, 1:1000) antibody was used to detect the neuronal silencer tetanus toxin light chain. The washing solution was PBS with 0.5% TritonX100, and blocking was performed with 10% heat inactivated goat serum. The brains were counterstained with anti-Bruchpilot (nc82, mouse, 1:20 dilution) and the anti-mouse Alexa633 secondary antibody.

To detect native expression of LOV-LexA (tdTomato) and CsChrimson:Venus in Figures 4C,D and S4, the flies were cold anesthetized after video recording and dissected immediately. The brains were fixed in 4% paraformaldehyde, counterstained with DAPI (1:3000 dilution in PBST), and imaged within two hours after mounting on a slide, as described previously (Ribeiro et al., 2022).

### Connectome analyses

We used the FlyWire and the NeuPrint interfaces to query the female (FAFB v783) and the male (MCNS v0.9) brain connectomes, respectively. The rood cell ID numbers as well as the number of synapses for each root cell ID pair were obtained from FlyWire and used to derive the number of root IDs and neuron types downstream of LC10a, LC10d neurons. The root cell IDs are provided in Table S2. The percentage of outputs of LC10d root id (Figure 5D) was calculated as the:

> (# synapses LC10d root ID --> the downstream root ID / total output synapses LC10d root ID) x 100

And averaged for each LC10d --> downstream neuron type. The percentage of total inputs to each descending neuron (DN) in Figure S4P was calculated similarly.

## Acknowledgements

We are grateful to A.H.Ali, C.Busch, C.Coskun, J.Haag, R.Kutlesa, B.M.Leonte, N.Pirogova, C. Theile, and B.Zuidinga for expertise help with various aspects of this work, and to all members of the A.Borst and M.Merrow groups for feedback. We thank Jürgen Haag and Martha Merrow for critically reading this manuscript, Dirk Reiff for sending *norpA* flies, and the FlyBank and FlyLight (Janelia Research Campus, HHMI) for sharing resources before publication. We are in debt to the FlyBase, the VDRC, and the Bloomington Drosophila Stock Center (NIH P40OD018537). This work was funded by the Max-Planck-Gesellschaft (N.D., M.S., S.P. and A.B.) and the Deutsche Forschungsgemeinschaft (DFG RI 3644/2-1), I.M.A.R. and C.W-Q..

## Author Contributions

I.M.A.R. designed all experiments with feedback from C.W-Q. and A.B.. C.W-Q. performed experiments and analysis presented in Figures S1H,I, S2G-J, 4A,B, 5A,D,E,F,J and S5O. The connectome analysis presented in figures 5B,C and S4M,N was performed by N.D.; dissections for figure S3J-N were performed by M.S.; the 2-floor assay chambers and previous iterations of them were designed and created by S.P., as was the light box used for optogenetic stimulation. I.M.A.R. performed all other experiments and wrote the paper with input from A.B., C.W-Q., N.D., and S.P..

---

**Figure S1:** Social interactions with expansion of sensory cues. **A.** Schematic showing the parameters orienting angle and distance. **B-E.** Median inter-fly distance (B), median absolute orienting angle male to female (C), percentage of time the male spends following (D) or with a single wing extended (E) for parental control (N=53), LC10s-SS2 block (N=28), LC10a-SS1 block (N=14), LC10bc-OL23B block (N=29), LC10d- SS3822 block (N=31), LC11-OL15B block (N=20), LC6-OL77B block (N=19), and LC9-SS2651 block (N=26). (**) p < 0.01, (***) p < 0.001, Student T test. **F-G.** Male-centered heatmap for difference after - before heat maps for randomly chosen points for the entire duration of the single pair assay (pre-courtship and courtship periods) for Canton S (N=32) and parental control (N=53). **H-K.** Male-centered heatmap for difference after - before heat maps for changes in forward velocity (accelerations and decelerations together) for *Canton S* (N=13), *norpA7-/-* (N=5), and for controls (N=53), LC10a-SS1-block (N=14), and LC10d-SS3822 block (N=31), for the pre-courtship (H, J) and courtship (I, K) periods. **L,M.** Amplitude of forward and rotational velocities (top), and forward and absolute rotational velocities profiles (bottom) for forward accelerations, decelerations and rotational change points, for the indicated genotypes. **N,O.** Male – male single pair assay for controls (N=8), and LC10d-SS3822 block (N=8). **N.** Male-centered heatmap of relative positions of the other male, for the 10-minute duration of the assay. **O.** Latency to the first occlusion (or close proximity, left) or the first wing extension (center), and percentage of time spent with wing extension (right). (*) p < 0.05, Student T test.

**Figure S2.** Female behavior (supplement to Figure 1) and avoidance in a vision-only assay (supplement to Figure 2). **A-B.** Forward and absolute rotational velocity for the time envelope around male forward accelerations (time point 0), for male (pink) and female (grey), for the indicated genotypes shown in Figure 1. **C.** Schematic showing orienting angles male to female and female to male. **D.** Change in absolute orienting angle male to female (pink) and female to male (grey) for the time envelope around male forward accelerations (time point 0). **E.** Schematic showing distance from the head of the one fly to the tip of the abdomen of the other fly, for male to female (pink) and female to male (grey). **F.** Change in distance head to abdomen male to female (pink) and female to male (grey) for the time envelope around male forward accelerations (time point 0). **G-N.** Male centered heatmap for difference after – before change points for all forward change points (G,I,KM) or randomly chosen points (H,J,L,N), for *Canton S* (N=62), and *norpA7-/-* (N=40), controls (N=16) and LC10d-SS3822 block (N=35). **O-Q.** Amplitude of forward and rotational velocities (left panels) and forward and absolute rotational velocities profiles (right panels) for forward accelerations (O), decelerations (P) and rotational change points (Q), for the indicated genotypes, controls (N=16), LC10a-SS1 block (N=26), and LC10d-SS3822 block (N=35), same as in Figure 2.

**Figure S3.** Analysis of collision avoidance (supplement to Figure 2) and morphological and functional characterization of LC10d neurons (supplement to Figure 3). **A-E.** Behavior analysis for collision points selected for the anterior visual field of the male (A) with interfly distances below 2mm, in single pair assays in large 42 mm chambers (B). **C-E.** Male walking speed (top row) and female angular size (bottom row) for the time leading up to a collision point for the pre-courtship period, for female relative motion back-to-front (regressive, brown or pink) and front-to-back (progressive, grey or pinkish grey), for the indicated genotypes, as in Figure 1. **F-I.** Behavior analysis for collision points selected for the anterior visual field of the male with interfly distances below 2mm, in vision-only, 2-floor assays (F). **G-I.** Male walking speed (top row) and female angular size (bottom row) for the time leading up to a collision point for female relative motion back-to-front (regressive, brown or pink) and front-to- back (progressive, grey or pinkish grey), for the indicated genotypes, as in Figure 2. **J.** Z-projection of a female brain expressing Venus under control of LC10a-LexA and tdTomato under control of LC10d-SS3822 driver. **K-N.** Genotypes: LC10a-LexA > CsChrimson:Venus, LC10a-SS1 > myrTomato (N = 6); LC10a-LexA > CsChrimson:Venus, LC10bc-OL23B > myrTomato (N = 2); LC10a-LexA > CsChrimson:Venus, LC10d-SS3822 > myrTomato (N = 7); **K.** Proportion of cells expressing only LC10a-LexA, or only a split-GAL4 driver either for LC10a, LC10bc and LC10d, or both. **L-N.** Proportion of tdTomato only (L), Venus only (M) or both (N) cells for each split driver in combination with LC10a-LexA. **O-Q.** Calcium responses plotted as normalized change in fluorescence over base line fluorescence (ΔF/F_0_) in response to edges in 8 directions bright or dark (P), full field flicker (Q) and counter-phase flicker (R).

**Figure S4:** LC10a and LC10d activity in vision-only or full cocktail of female sensory cues and optogenetic activation of a subset of LC10a or LC10d neurons. **A-D.** Schematic showing vision-only, 2-floors assay (A) and full cocktail of female sensory cues (C), along with the distribution of absolute orienting angle male to female and inter-fly distance (B,D). **E,F.** Schematic showing CRTC:EGFP localization in the nucleus in the presence of neuronal activity (E) and the odds ratio of CRTC:EGFP/DAPi and CRTC:EGFP/mCherry as a proxy for localization in the nucleus (F). (*) p < 0.05, Student T test. **B,D,F.** Genotypes: LC10a-SS1 > CRTC:EGFP without female (N=4) LC10a-SS1 > CRTC:EGFP with female (N=2) LC10d-SS3822 > CRTC:EGFP without female (N=4) LC10d-SS3822 > CRTC:EGFP with female (N=4) **G-L.** Light-gated unilateral expression of CsChrimson:Venus in 2 to 7 cells in LC10a-SS1 (G-I), and LC10d-SS3822 (J-L), with LC10a-SS1 > LexA-LOV, CsChrimsonVenus (N=12, G), and LC10d-SS3822 > LexA-LOV, CsChrimsonVenus (N=16, J). **G,J.** Z-projections of brains of representative unilateral expression, merge on the left and Venus on the right. **H,K.** Relative turning over time, with red shade indicating the time the red-light stimulus was on. **I,L.** Walking speed over time, with red shade indicating the time the red-light stimulus was on. **M-N.** Flywire representation of all LC10d neurons that are upstream the left DNa10 neuron (M) and in a planar view of the lobula (N). **O.** Number of synapses for LC10a or LC10d cells with 5 or more synapses onto the ipsilateral DNa10, for female (left) and male (right). (*) p < 0.05, (***) p < 0.001, Student T-test. **P.** Mean number of synapses per cell ID to descending neurons from visual projection neurons upstream DNa10 and other descending neurons from the male connectome.

**Figure S5:** LC10d-DNa10 mediated avoidance, supplement to Figure 5. **A-D.** Behavior analysis for collision points selected for the anterior visual field of the male (A) with interfly distances below 2mm, in vision-only, 2-floor assays (B). **C-F.** Male walking speed (top row) and female angular size (bottom row) for the time leading up to a collision point for female relative motion back-to-front (regressive, brown or pink) and front-to- back (progressive, grey or pinkish grey), for the indicated genotypes, as in Figure 5G-I. **G-L.** Behavior analysis for collision points selected for the anterior visual field of the male in single pair assays with the female as test fly (G) with interfly distances below 2mm. **H-L.** Female walking speed (top row) and male angular size (bottom row) for the time leading up to a collision point for male relative motion back-to-front (regressive, brown or blue) and front-to-back (progressive, grey or blueish grey), for the indicated genotypes, as in Figure 5K-M. **M,N.** Latency to male courtship initiation with first wing extension (M) and following (N) for females with the genotypes: parental control (N=15); LC10d-SS3822 block (N=14); DNa10/04-SS2384 block (N=14); parental control 2 (N=4); and parental control 3 (N=15). **O.** Projection of the male relative positions along the female visual anterior-posterior axis resulting from the difference after-before for female forward accelerations. Mean A.U.C. for the anterior visual field with p > 0.05 for female control versus DNa10/04-SS2384 block, Student T-test. Genotypes as in Figure 5M. **P-R.** Amplitude of female forward and rotational velocities (left panels) and forward and absolute rotational velocities profiles (right panels) for forward velocity accelerations (M), decelerations (N) and rotational velocity change points (O), for the indicated genotypes, as in Figure 5K-M.

## References

Ache JM, Polsky J, Alghailani S, Parekh R, Breads P, Peek MY, Bock DD, Von Reyn CR, Card GM. 2019. Neural Basis for Looming Size and Velocity Encoding in the Drosophila Giant Fiber Escape Pathway. Current Biology 29:1073–1081.e4. DOI: 10.1016/j.cub.2019.01.079

Agrawal S, Safarik S, Dickinson M. 2014. The relative roles of vision and chemosensation in mate recognition of Drosophila melanogaster. J Exp Biol 217:2796–805. DOI: 10.1242/jeb.105817

Alekseyenko OV, Chan Y-B, Okaty BW, Chang Y, Dymecki SM, Kravitz EA. 2019. Serotonergic Modulation of Aggression in Drosophila Involves GABAergic and Cholinergic Opposing Pathways. Current Biology 29:2145–2156.e5. DOI: 10.1016/j.cub.2019.05.070

Allen KM, Lawlor J, Salles A, Moss CF. 2021. Orienting our view of the superior colliculus: specializations and general functions. Current Opinion in Neurobiology 71:119–126. DOI: 10.1016/j.conb.2021.10.005

Aptekar JW, Shoemaker PA, Frye MA. 2012. Figure Tracking by Flies Is Supported by Parallel Visual Streams. Current Biology 22:482–487. DOI: 10.1016/j.cub.2012.01.044

Arenz A, Drews MS, Richter FG, Ammer G, Borst A. 2017. The Temporal Tuning of the Drosophila Motion Detectors Is Determined by the Dynamics of Their Input Elements. Current Biology 27:929–944. DOI: 10.1016/j.cub.2017.01.051

Bahl A, Ammer G, Schilling T, Borst A. 2013. Object tracking in motion-blind flies. Nature Neuroscience 16:730– 738. DOI: 10.1038/nn.3386

Bath DE, Stowers JR, Hormann D, Poehlmann A, Dickson BJ, Straw AD. 2014. FlyMAD: rapid thermogenetic control of neuronal activity in freely walking Drosophila. Nat Methods 11:756–62. DOI: 10.1038/nmeth.2973

Berg S, Beckett IR, Costa M, Schlegel P, Januszewski M, Marin EC, Nern A, Preibisch S, Qiu W, Takemura Shinya, Fragniere AMC, Champion AS, Adjavon D-Y, Cook M, Gkantia M, Hayworth KJ, Huang GB, Kampf F, Katz WT, Lu Z, Ordish C, Paterson T, Stuerner T, Trautman ET, Whittle CR, Burnett LE, Hoeller J, Li F, Loesche F, Morris BJ, Pietzsch T, Pleijzier MW, Silva V, Yin Y, Ali I, Badalamente G, Bates AS, Bogovic J, Brooks P, Cachero S, Canino BS, Chaisrisawatsuk B, Clements J, Crowe A, De Haan Vicente I, Dempsey G, Dona E, Dos Santos M, Dreher M, Dunne CR, Eichler K, Finley-May S, Flynn MA, Hameed I, Hopkins GP, Hubbard PM, Kiassat L, Kovalyak J, Lauchie SA, Leonard M, Lohff A, Longden KD, Maldonado CA, Mitletton M, Moitra I, Moon SS, Mooney C, Munnelly EJ, Okeoma N, Olbris DJ, Pai A, Patel B, Phillips EM, Plaza SM, Richards A, Rivas Salinas J, Roberts RJV, Rogers EM, Scott AL, Scuderi LA, Seenivasan P, Serratosa Capdevila L, Smith C, Svirskas R, Takemura Satoko, Tastekin I, Thomson A, Umayam L, Walsh JJ, Whittome H, Xu CS, Yakal EA, Yang T, Zhao A, George R, Jain V, Jayaraman V, Korff W, Meissner GW, Romani S, Funke J, Knecht C, Saalfeld S, Scheffer LK, Waddell S, Card GM, Ribeiro C, Reiser MB, Hess HF, Rubin GM, Jefferis GSXE. 2025. Sexual dimorphism in the complete connectome of the Drosophila male central nervous system. DOI: 10.1101/2025.10.09.680999

Bertsch DJ, Palacios Castillo LM, Frye MA. 2025. Serotonin selectively modulates visual responses of object motion detectors in *Drosophila*. Journal of Neurophysiology 134:962–984. DOI: 10.1152/jn.00154.2025

Bidaye SS, Laturney M, Chang AK, Liu Y, Bockemühl T, Büschges A, Scott K. 2020. Two Brain Pathways Initiate Distinct Forward Walking Programs in Drosophila. Neuron 108:469–485.e8. DOI: 10.1016/j.neuron.2020.07.032

Bonheur M, Swartz KJ, Metcalf MG, Wen X, Zhukovskaya A, Mehta A, Connors KE, Barasch JG, Jamieson AR, Martin KC, Axel R, Hattori D. 2023. A rapid and bidirectional reporter of neural activity reveals neural correlates of social behaviors in Drosophila. Nature Neuroscience 26:1295–1307. DOI: 10.1038/s41593-023-01357-w

Borst A, Groschner LN. 2023. How Flies See Motion. Annual Review of Neuroscience 46:17–37. DOI: 10.1146/annurev-neuro-080422-111929

Briceno RD. 2003. Sexual behavior of mutant strains of the medfly Ceratitis capitata (Diptera: Tephritidae). Rev Biol Trop 51:763–7.

Budick SA, O’Malley DM. 2000. Locomotor Repertoire of The Larval Zebrafish: Swimming, Turning and Prey Capture. Journal of Experimental Biology 203:2565–2579. DOI: 10.1242/jeb.203.17.2565

Chen T-W, Wardill TJ, Sun Y, Pulver SR, Renninger SL, Baohan A, Schreiter ER, Kerr RA, Orger MB, Jayaraman V, Looger LL, Svoboda K, Kim DS. 2013. Ultrasensitive fluorescent proteins for imaging neuronal activity. Nature 499:295–300. DOI: 10.1038/nature12354

Cheng KY, Colbath RA, Frye MA. 2019. Olfactory and Neuromodulatory Signals Reverse Visual Object Avoidance to Approach in Drosophila. Curr Biol 29:2058–2065 e2. DOI: 10.1016/j.cub.2019.05.010

Cheong HS, Eichler K, Stürner T, Asinof SK, Champion AS, Marin EC, Oram TB, Sumathipala M, Venkatasubramanian L, Namiki S, Siwanowicz I, Costa M, Berg S, FlyEM Project Team J, Jefferis GS, Card GM. 2026. Organization of circuits linking descending input to motor output in the Drosophila Male Adult Nerve Cord connectome. eLife 13:RP96084. DOI: 10.7554/eLife.96084.3

Cheong HS, Siwanowicz I, Card GM. 2020. Multi-regional circuits underlying visually guided decision-making in Drosophila. Curr Opin Neurobiol 65:77–87. DOI: 10.1016/j.conb.2020.10.010

Clowney EJ, Iguchi S, Bussell JJ, Scheer E, Ruta V. 2015. Multimodal Chemosensory Circuits Controlling Male Courtship in Drosophila. Neuron 87:1036–49. DOI: 10.1016/j.neuron.2015.07.025

Coen P, Clemens J, Weinstein AJ, Pacheco DA, Deng Y, Murthy M. 2014. Dynamic sensory cues shape song structure in Drosophila. Nature 507:233–237. DOI: 10.1038/nature13131

Coen P, Murthy M. 2016. Singing on the fly: sensorimotor integration and acoustic communication in Drosophila. Current Opinion in Neurobiology 38:38–45. DOI: 10.1016/j.conb.2016.01.013

Collie MF, Jin C, Rockwell V, Kellogg E, Vanderbeck QX, Hartman AK, Holtz SL, Wilson RI. 2026. Specialized parallel pathways for adaptive control of visual object pursuit. Neuron S0896627326000012. DOI: 10.1016/j.neuron.2026.01.001

Cook R. 1980. The extent of visual control in the courtship tracking of D. melanogaster. Biological Cybernetics 37:41–51. DOI: 10.1007/BF00347641

Cook R. 1979. The courtship tracking of Drosophila melanogaster. Biological Cybernetics 34:91–106. DOI: 10.1007/BF00365473

Cowley BR, Calhoun AJ, Rangarajan N, Ireland E, Turner MH, Pillow JW, Murthy M. 2024. Mapping model units to visual neurons reveals population code for social behaviour. Nature 629:1100–1108. DOI: 10.1038/s41586-024-07451-8

Dorkenwald S, Matsliah A, Sterling AR, Schlegel P, Yu SC, McKellar CE, Lin A, Costa M, Eichler K, Yin Y, Silversmith W, Schneider-Mizell C, Jordan CS, Brittain D, Halageri A, Kuehner K, Ogedengbe O, Morey R, Gager J, Kruk K, Perlman E, Yang R, Deutsch D, Bland D, Sorek M, Lu R, Macrina T, Lee K, Bae JA, Mu S, Nehoran B, Mitchell E, Popovych S, Wu J, Jia Z, Castro MA, Kemnitz N, Ih D, Bates AS, Eckstein N, Funke J, Collman F, Bock DD, Jefferis G, Seung HS, Murthy M, FlyWire C. 2024. Neuronal wiring diagram of an adult brain. Nature 634:124–138. DOI: 10.1038/s41586-024-07558-y

Duistermars BJ, Pfeiffer BD, Hoopfer ED, Anderson DJ. 2018. A Brain Module for Scalable Control of Complex, Multi-motor Threat Displays. Neuron 100:1474–1490.e4. DOI: 10.1016/j.neuron.2018.10.027, PMID: 30415997

Dweck HK, Ebrahim SA, Thoma M, Mohamed AA, Keesey IW, Trona F, Lavista-Llanos S, Svatos A, Sachse S, Knaden M, Hansson BS. 2015. Pheromones mediating copulation and attraction in Drosophila. Proc Natl Acad Sci U S A 112:E2829–35. DOI: 10.1073/pnas.1504527112

Ferreira CH, Moita MA. 2020. Behavioral and neuronal underpinnings of safety in numbers in fruit flies. Nature Communications 11:4182. DOI: 10.1038/s41467-020-17856-4, PMID: 32826882

Fischbach K-F, Dittrich APM. 1989. The optic lobe of Drosophila melanogaster. I. A Golgi analysis of wild-type structure. Cell and Tissue Research 258. DOI: 10.1007/BF00218858

Hall JC. 1994. The Mating of a Fly. Science 264:1702–1714. DOI: 10.1126/science.8209251

Helmbrecht TO, Dal Maschio M, Donovan JC, Koutsouli S, Baier H. 2018. Topography of a Visuomotor Transformation. Neuron 100:1429–1445.e4. DOI: 10.1016/j.neuron.2018.10.021

Hindmarsh Sten T, Li R, Otopalik A, Ruta V. 2021. Sexual arousal gates visual processing during Drosophila courtship. Nature 595:549–553. DOI: 10.1038/s41586-021-03714-w

Hoeller J, Zhao A, Nern A, Rogers EM, Romani S, Reiser MB. 2025. The organization of visual pathways in the Drosophila brain. DOI: 10.64898/2025.12.22.696097

Inagaki HK, Jung Y, Hoopfer ED, Wong AM, Mishra N, Lin JY, Tsien RY, Anderson DJ. 2014. Optogenetic control of Drosophila using a red-shifted channelrhodopsin reveals experience-dependent influences on courtship. Nat Methods 11:325–32. DOI: 10.1038/nmeth.2765

Isa T, Marquez-Legorreta E, Grillner S, Scott EK. 2021. The tectum/superior colliculus as the vertebrate solution for spatial sensory integration and action. Current Biology 31:R741–R762. DOI: 10.1016/j.cub.2021.04.001

Kappel JM, Förster D, Slangewal K, Shainer I, Svara F, Donovan JC, Sherman S, Januszewski M, Baier H, Larsch J. 2022. Visual recognition of social signals by a tectothalamic neural circuit. Nature 608:146–152. DOI: 10.1038/s41586-022-04925-5, PMID: 35831500

Keleş MF, Frye MA. 2017. Object-Detecting Neurons in Drosophila. Current Biology 27:680–687. DOI: 10.1016/j.cub.2017.01.012

Klapoetke NC, Murata Y, Kim SS, Pulver SR, Birdsey-Benson A, Cho YK, Morimoto TK, Chuong AS, Carpenter EJ, Tian Z, Wang J, Xie Y, Yan Z, Zhang Y, Chow BY, Surek B, Melkonian M, Jayaraman V, Constantine-Paton M, Wong GK-S, Boyden ES. 2014. Independent optical excitation of distinct neural populations. Nature Methods 11:338–346. DOI: 10.1038/nmeth.2836

Klapoetke NC, Nern A, Rogers EM, Rubin GM, Reiser MB, Card GM. 2022. A functionally ordered visual feature map in the Drosophila brain. Neuron 110:1700–1711.e6. DOI: 10.1016/j.neuron.2022.02.013

Koh T-W, He Z, Gorur-Shandilya S, Menuz K, Larter NK, Stewart S, Carlson JR. 2014. The Drosophila IR20a Clade of Ionotropic Receptors Are Candidate Taste and Pheromone Receptors. Neuron 83:850–865. DOI: 10.1016/j.neuron.2014.07.012

Kohatsu S, Yamamoto D. 2015. Visually induced initiation of Drosophila innate courtship-like following pursuit is mediated by central excitatory state. Nature Communications 6:6457. DOI: 10.1038/ncomms7457

Land MF, Collett TS. 1974. Chasing behaviour of houseflies (Fannia canicularis): A description and analysis. Journal of Comparative Physiology 89:331–357. DOI: 10.1007/BF00695351

Larsch J, Baier H. 2018. Biological Motion as an Innate Perceptual Mechanism Driving Social Affiliation. Current biology: CB 28:3523–3532.e4. DOI: 10.1016/j.cub.2018.09.014, PMID: 30393036

Li XL, Li DD, Cai XY, Cheng DF, Lu YY. 2024. Reproductive behavior of fruit flies: courtship, mating, and oviposition. Pest Manag Sci 80:935–952. DOI: 10.1002/ps.7816

Lu B, LaMora A, Sun Y, Welsh MJ, Ben-Shahar Y. 2012. ppk23-Dependent chemosensory functions contribute to courtship behavior in Drosophila melanogaster. PLoS Genet 8:e1002587. DOI: 10.1371/journal.pgen.1002587

Maimon G, Straw AD, Dickinson MH. 2008. A Simple Vision-Based Algorithm for Decision Making in Flying Drosophila. Current Biology 18:464–470. DOI: 10.1016/j.cub.2008.02.054

Maisak MS, Haag J, Ammer G, Serbe E, Meier M, Leonhardt A, Schilling T, Bahl A, Rubin GM, Nern A, Dickson BJ, Reiff DF, Hopp E, Borst A. 2013. A directional tuning map of Drosophila elementary motion detectors. Nature 500:212–216. DOI: 10.1038/nature12320

Markow TA. 2015. The secret lives of Drosophila flies. eLife 4:e06793. DOI: 10.7554/eLife.06793

Markow TA. 1987. Behavioral and Sensory Basis of Courtship Success in Drosophila melanogaster. Proceedings of the National Academy of Sciences of the United States of America 84:6200–6204.

Matsliah A, Yu S, Kruk K, Bland D, Burke AT, Gager J, Hebditch J, Silverman B, Willie KP, Willie R, Sorek M, Sterling AR, Kind E, Garner D, Sancer G, Wernet MF, Kim SS, Murthy M, Seung HS, The FlyWire Consortium, David C, Joroff J, Kristiansen A, Stocks T, Braun A, Silies M, Skelton J, Aiken TR, Ioannidou M, Collie M, Linneweber GA, Molina-Obando S, Dorkenwald S, Panes N, Gogo AM, Rastgarmoghaddam D, Pilapil C, Candilada RA, Serafetinidis N, Lee W-C, Borst A, Wilson RI, Schlegel P, Jefferis GSXE. 2024. Neuronal parts list and wiring diagram for a visual system. Nature 634:166–180. DOI: 10.1038/s41586-024-07981-1

Mauss AS, Meier M, Serbe E, Borst A. 2014. Optogenetic and Pharmacologic Dissection of Feedforward Inhibition in *Drosophila* Motion Vision. The Journal of Neuroscience 34:2254–2263. DOI: 10.1523/JNEUROSCI.3938-13.2014

McKinney RM, Ben-Shahar Y. 2019. Visual recognition of the female body axis drives spatial elements of male courtship in Drosophila melanogaster. DOI: 10.1101/576322

Meissner GW, Vannan A, Jeter J, Close K, DePasquale GM, Dorman Z, Forster K, Beringer JA, Gibney TV, Hausenfluck JH, He Y, Henderson K, Johnson L, Johnston RM, Ihrke G, Iyer N, Lazarus R, Lee K, Li H-H, Liaw H-P, Melton B, Miller S, Motaher R, Novak A, Ogundeyi O, Petruncio A, Price J, Protopapas S, Tae S, Taylor J, Vorimo R, Yarbrough B, Zeng KX, Zugates CT, Dionne H, Angstadt C, Ashley K, Cavallaro A, Dang T, Gonzalez GA, Hibbard KL, Huang C, Kao J-C, Laverty T, Mercer M, Perez B, Pitts S, Ruiz D, Vallanadu V, Zheng GZ, Goina C, Otsuna H, Rokicki K, Svirskas RR, Cheong HSJ, Dolan M-J, Ehrhardt E, Feng K, El Galfi B, Goldammer J, Huston SJ, Hu N, Ito M, McKellar C, Minegishi R, Namiki S, Nern A, Schretter CE, Sterne GR, Venkatasubramanian L, Wang K, Wolff T, Wu M, George R, Malkesman O, Aso Y, Card GM, Dickson BJ, Korff W, Ito K, Truman JW, Zlatic M, Rubin GM. 2024. A split-GAL4 driver line resource for Drosophila CNS cell types.

Monavarfeshani A, Sabbagh U, Fox MA. 2017. Not a one-trick pony: Diverse connectivity and functions of the rodent lateral geniculate complex. Visual Neuroscience 34:E012. DOI: 10.1017/S0952523817000098

Namiki S, Dickinson MH, Wong AM, Korff W, Card GM. 2018. The functional organization of descending sensory-motor pathways in Drosophila. eLife 7:e34272. DOI: 10.7554/eLife.34272

Nern A, Loesche F, Takemura Shin-ya, Burnett LE, Dreher M, Gruntman E, Hoeller J, Huang GB, Januszewski M, Klapoetke NC, Koskela S, Longden KD, Lu Z, Preibisch S, Qiu W, Rogers EM, Seenivasan P, Zhao A, Bogovic J, Canino BS, Clements J, Cook M, Finley-May S, Flynn MA, Hameed I, Fragniere AMC, Hayworth KJ, Hopkins GP, Hubbard PM, Katz WT, Kovalyak J, Lauchie SA, Leonard M, Lohff A, Maldonado CA, Mooney C, Okeoma N, Olbris DJ, Ordish C, Paterson T, Phillips EM, Pietzsch T, Salinas JR, Rivlin PK, Schlegel P, Scott AL, Scuderi LA, Takemura Satoko, Talebi I, Thomson A, Trautman ET, Umayam L, Walsh C, Walsh JJ, Xu CS, Yakal EA, Yang T, Zhao T, Funke J, George R, Hess HF, Jefferis GSXE, Knecht C, Korff W, Plaza SM, Romani S, Saalfeld S, Scheffer LK, Berg S, Rubin GM, Reiser MB. 2025. Connectome-driven neural inventory of a complete visual system. Nature 641:1225–1237. DOI: 10.1038/s41586-025-08746-0

Nikonov SS, Kholodenko R, Lem J, Pugh EN. 2006. Physiological Features of the S- and M-cone Photoreceptors of Wild-type Mice from Single-cell Recordings. The Journal of General Physiology 127:359–374. DOI: 10.1085/jgp.200609490

Nojima T, Rings A, Allen AM, Otto N, Verschut TA, Billeter J-C, Neville MC, Goodwin SF. 2021. A sex-specific switch between visual and olfactory inputs underlies adaptive sex differences in behavior. Current Biology 31:1175–1191.e6. DOI: 10.1016/j.cub.2020.12.047

Nordström K, Barnett PD, Moyer De Miguel IM, Brinkworth RSA, O’Carroll DC. 2008. Sexual Dimorphism in the Hoverfly Motion Vision Pathway. Current Biology 18:661–667. DOI: 10.1016/j.cub.2008.03.061

Nordstrom K, Barnett PD, O’Carroll DC. 2006. Insect detection of small targets moving in visual clutter. PLoS Biol 4:e54. DOI: 10.1371/journal.pbio.0040054

Olberg RM. 2012. Visual control of prey-capture flight in dragonflies. Current Opinion in Neurobiology 22:267– 271. DOI: 10.1016/j.conb.2011.11.015

Otsuna H, Ito K. 2006. Systematic analysis of the visual projection neurons of Drosophila melanogaster. I. Lobula-specific pathways. J Comp Neurol 497:928–58. DOI: 10.1002/cne.21015

Pan Y, Meissner GW, Baker BS. 2012. Joint control of *Drosophila* male courtship behavior by motion cues and activation of male-specific P1 neurons. Proceedings of the National Academy of Sciences 109:10065– 10070. DOI: 10.1073/pnas.1207107109

Plaza SM, Clements J, Dolafi T, Umayam L, Neubarth NN, Scheffer LK, Berg S. 2022. neuPrint: An open access tool for EM connectomics. Frontiers in Neuroinformatics 16:896292. DOI: 10.3389/fninf.2022.896292

Portugues R, Feierstein CE, Engert F, Orger MB. 2014. Whole-Brain Activity Maps Reveal Stereotyped, Distributed Networks for Visuomotor Behavior. Neuron 81:1328–1343. DOI: 10.1016/j.neuron.2014.01.019

Rayshubskiy A, Holtz SL, Bates AS, Vanderbeck QX, Serratosa Capdevila L, Rockwell V, Wilson R. 2025. Neural circuit mechanisms for steering control in walking Drosophila. eLife 13:RP102230. DOI: 10.7554/eLife.102230.3

Ribeiro IMA, Drews M, Bahl A, Machacek C, Borst A, Dickson BJ. 2018. Visual Projection Neurons Mediating Directed Courtship in Drosophila. Cell 174:607–621.e18. DOI: 10.1016/j.cell.2018.06.020

Ribeiro IMA, Essbauer W, Kutlesa R, Borst A. 2022. Spatial and temporal control of expression with light-gated LOV-LexA. G3 (Bethesda) 12. DOI: 10.1093/g3journal/jkac178

Rings A, Goodwin SF. 2019. To court or not to court - a multimodal sensory decision in Drosophila males. Curr Opin Insect Sci 35:48–53. DOI: 10.1016/j.cois.2019.06.009

Rubin GM, Managan C, Dreher M, Kim E, Miller S, Boone K, Robie AA, Taylor AL, Branson K, Schretter CE, Otopalik AG. 2025. Networks of sexually dimorphic neurons that regulate social behaviors in Drosophila. DOI: 10.1101/2025.10.21.683766

Sayeed O, Benzer S. 1996. Behavioral genetics of thermosensation and hygrosensation in Drosophila. Proc Natl Acad Sci U S A 93:6079–84. DOI: 10.1073/pnas.93.12.6079

Scheffer LK, Xu CS, Januszewski M, Lu Z, Takemura SY, Hayworth KJ, Huang GB, Shinomiya K, Maitlin-Shepard J, Berg S, Clements J, Hubbard PM, Katz WT, Umayam L, Zhao T, Ackerman D, Blakely T, Bogovic J, Dolafi T, Kainmueller D, Kawase T, Khairy KA, Leavitt L, Li PH, Lindsey L, Neubarth N, Olbris DJ, Otsuna H, Trautman ET, Ito M, Bates AS, Goldammer J, Wolff T, Svirskas R, Schlegel P, Neace E, Knecht CJ, Alvarado CX, Bailey DA, Ballinger S, Borycz JA, Canino BS, Cheatham N, Cook M, Dreher M, Duclos O, Eubanks B, Fairbanks K, Finley S, Forknall N, Francis A, Hopkins GP, Joyce EM, Kim S, Kirk NA, Kovalyak J, Lauchie SA, Lohff A, Maldonado C, Manley EA, McLin S, Mooney C, Ndama M, Ogundeyi O, Okeoma N, Ordish C, Padilla N, Patrick CM, Paterson T, Phillips EE, Phillips EM, Rampally N, Ribeiro C, Robertson MK, Rymer JT, Ryan SM, Sammons M, Scott AK, Scott AL, Shinomiya A, Smith C, Smith K, Smith NL, Sobeski MA, Suleiman A, Swift J, Takemura S, Talebi I, Tarnogorska D, Tenshaw E, Tokhi T, Walsh JJ, Yang T, Horne JA, Li F, Parekh R, Rivlin PK, Jayaraman V, Costa M, Jefferis GS, Ito K, Saalfeld S, George R, Meinertzhagen IA, Rubin GM, Hess HF, Jain V, Plaza SM. 2020. A connectome and analysis of the adult Drosophila central brain. Elife 9. DOI: 10.7554/eLife.57443

Schlegel P, Yin Y, Bates AS, Dorkenwald S, Eichler K, Brooks P, Han DS, Gkantia M, Dos Santos M, Munnelly EJ, Badalamente G, Serratosa Capdevila L, Sane VA, Fragniere AMC, Kiassat L, Pleijzier MW, Sturner T, Tamimi IFM, Dunne CR, Salgarella I, Javier A, Fang S, Perlman E, Kazimiers T, Jagannathan SR, Matsliah A, Sterling AR, Yu SC, McKellar CE, FlyWire C, Costa M, Seung HS, Murthy M, Hartenstein V, Bock DD, Jefferis G. 2024. Whole-brain annotation and multi-connectome cell typing of Drosophila. Nature 634:139–152. DOI: 10.1038/s41586-024-07686-5

Schretter CE, Hindmarsh Sten T, Klapoetke N, Shao M, Nern A, Dreher M, Bushey D, Robie AA, Taylor AL, Branson K, Otopalik A, Ruta V, Rubin GM. 2025. Social state alters vision using three circuit mechanisms in Drosophila. Nature 637:646–653. DOI: 10.1038/s41586-024-08255-6

Semmelhack JL, Donovan JC, Thiele TR, Kuehn E, Laurell E, Baier H. 2014. A dedicated visual pathway for prey detection in larval zebrafish. eLife 3:e04878. DOI: 10.7554/eLife.04878

Seung HS. 2024. Predicting visual function by interpreting a neuronal wiring diagram. Nature 634:113–123. DOI: 10.1038/s41586-024-07953-5

Städele C, Keleş MF, Mongeau J-M, Frye MA. 2020. Non-canonical Receptive Field Properties and Neuromodulation of Feature-Detecting Neurons in Flies. Current Biology 30:2508–2519.e6. DOI: 10.1016/j.cub.2020.04.069

Strausfeld NJ. 2012. Arthropod Brains: Evolution, Functional Elegance, and Historical Significance. Harvard University Press. DOI: 10.2307/j.ctv1dp0v2h

Sweeney ST, Broadie K, Keane J, Niemann H, O’Kane CJ. 1995. Targeted expression of tetanus toxin light chain in Drosophila specifically eliminates synaptic transmission and causes behavioral defects. Neuron 14:341–351. DOI: 10.1016/0896-6273(95)90290-2

Tanaka R, Clark DA. 2022. Neural mechanisms to exploit positional geometry for collision avoidance. Current Biology 32:2357–2374.e6. DOI: 10.1016/j.cub.2022.04.023

Tanaka R, Clark DA. 2020. Object-Displacement-Sensitive Visual Neurons Drive Freezing in Drosophila. Current Biology 30:2532–2550.e8. DOI: 10.1016/j.cub.2020.04.068

Thistle R, Cameron P, Ghorayshi A, Dennison L, Scott K. 2012. Contact chemoreceptors mediate male-male repulsion and male-female attraction during Drosophila courtship. Cell 149:1140–51. DOI: 10.1016/j.cell.2012.03.045

Thyselius M, Gonzalez-Bellido PT, Wardill TJ, Nordström K. 2018. Visual approach computation in feeding hoverflies. Journal of Experimental Biology 221:jeb177162. DOI: 10.1242/jeb.177162

Toda H, Zhao X, Dickson BJ. 2012. The Drosophila female aphrodisiac pheromone activates ppk23(+) sensory neurons to elicit male courtship behavior. Cell Rep 1:599–607. DOI: 10.1016/j.celrep.2012.05.007

Von Reyn CR, Breads P, Peek MY, Zheng GZ, Williamson WR, Yee AL, Leonardo A, Card GM. 2014. A spike- timing mechanism for action selection. Nature Neuroscience 17:962–970. DOI: 10.1038/nn.3741

Von Reyn CR, Nern A, Williamson WR, Breads P, Wu M, Namiki S, Card GM. 2017. Feature Integration Drives Probabilistic Behavior in the Drosophila Escape Response. Neuron 94:1190–1204.e6. DOI: 10.1016/j.neuron.2017.05.036

Vrontou E, Nilsen SP, Demir E, Kravitz EA, Dickson BJ. 2006. fruitless regulates aggression and dominance in Drosophila. Nature Neuroscience 9:1469–1471. DOI: 10.1038/nn1809

Wiltschko AB, Johnson MJ, Iurilli G, Peterson RE, Katon JM, Pashkovski SL, Abraira VE, Adams RP, Datta SR. 2015. Mapping Sub-Second Structure in Mouse Behavior. Neuron 88:1121–1135. DOI: 10.1016/j.neuron.2015.11.031

Wu M, Nern A, Williamson WR, Morimoto MM, Reiser MB, Card GM, Rubin GM. 2016. Visual projection neurons in the Drosophila lobula link feature detection to distinct behavioral programs. Elife 5. DOI: 10.7554/eLife.21022

Yamamoto D, Koganezawa M. 2013. Genes and circuits of courtship behaviour in Drosophila males. Nature Reviews Neuroscience 14:681–692. DOI: 10.1038/nrn3567

Yu JY, Kanai MI, Demir E, Jefferis GSXE, Dickson BJ. 2010. Cellular Organization of the Neural Circuit that Drives Drosophila Courtship Behavior. Current Biology 20:1602–1614. DOI: 10.1016/j.cub.2010.08.025

Zabala F, Polidoro P, Robie A, Branson K, Perona P, Dickinson MH. 2012. A Simple Strategy for Detecting Moving Objects during Locomotion Revealed by Animal-Robot Interactions. Current Biology 22:1344– 1350. DOI: 10.1016/j.cub.2012.05.024

Zhao A, Gruntman E, Nern A, Iyer N, Rogers EM, Koskela S, Siwanowicz I, Dreher M, Flynn MA, Laughland C, Ludwig H, Thomson A, Moran C, Gezahegn B, Bock DD, Reiser MB. 2025. Eye structure shapes neuron function in Drosophila motion vision. Nature. DOI: 10.1038/s41586-025-09276-5

Zheng Z, Lauritzen JS, Perlman E, Robinson CG, Nichols M, Milkie D, Torrens O, Price J, Fisher CB, Sharifi N, Calle-Schuler SA, Kmecova L, Ali IJ, Karsh B, Trautman ET, Bogovic JA, Hanslovsky P, Jefferis G, Kazhdan M, Khairy K, Saalfeld S, Fetter RD, Bock DD. 2018. A Complete Electron Microscopy Volume of the Brain of Adult Drosophila melanogaster. Cell 174:730–743 e22. DOI: 10.1016/j.cell.2018.06.019

