## Supplementary figures and images for "A multi-input optic glomerulus mediates opposing behavioral responses to visual objects"

### Supplemental Figures

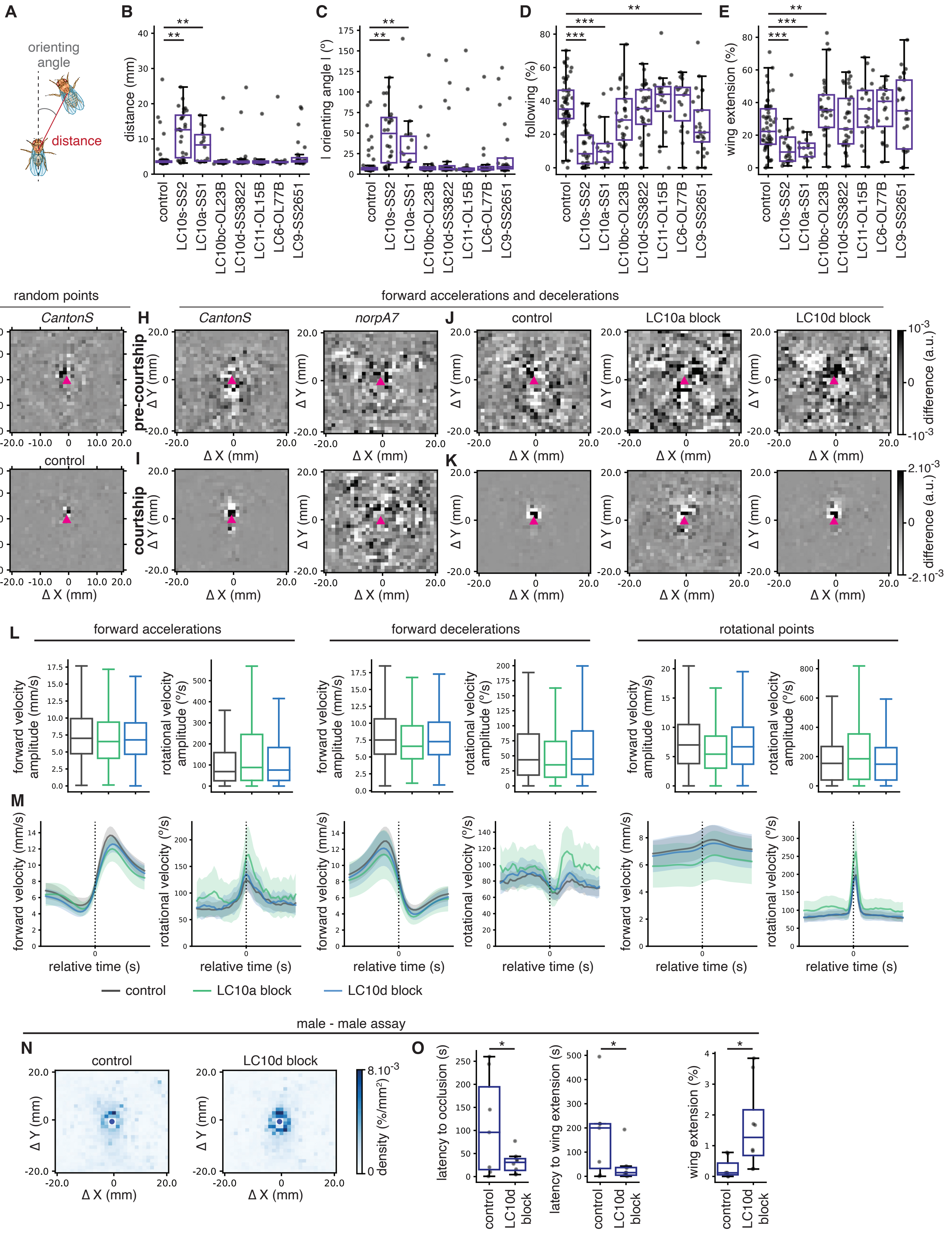

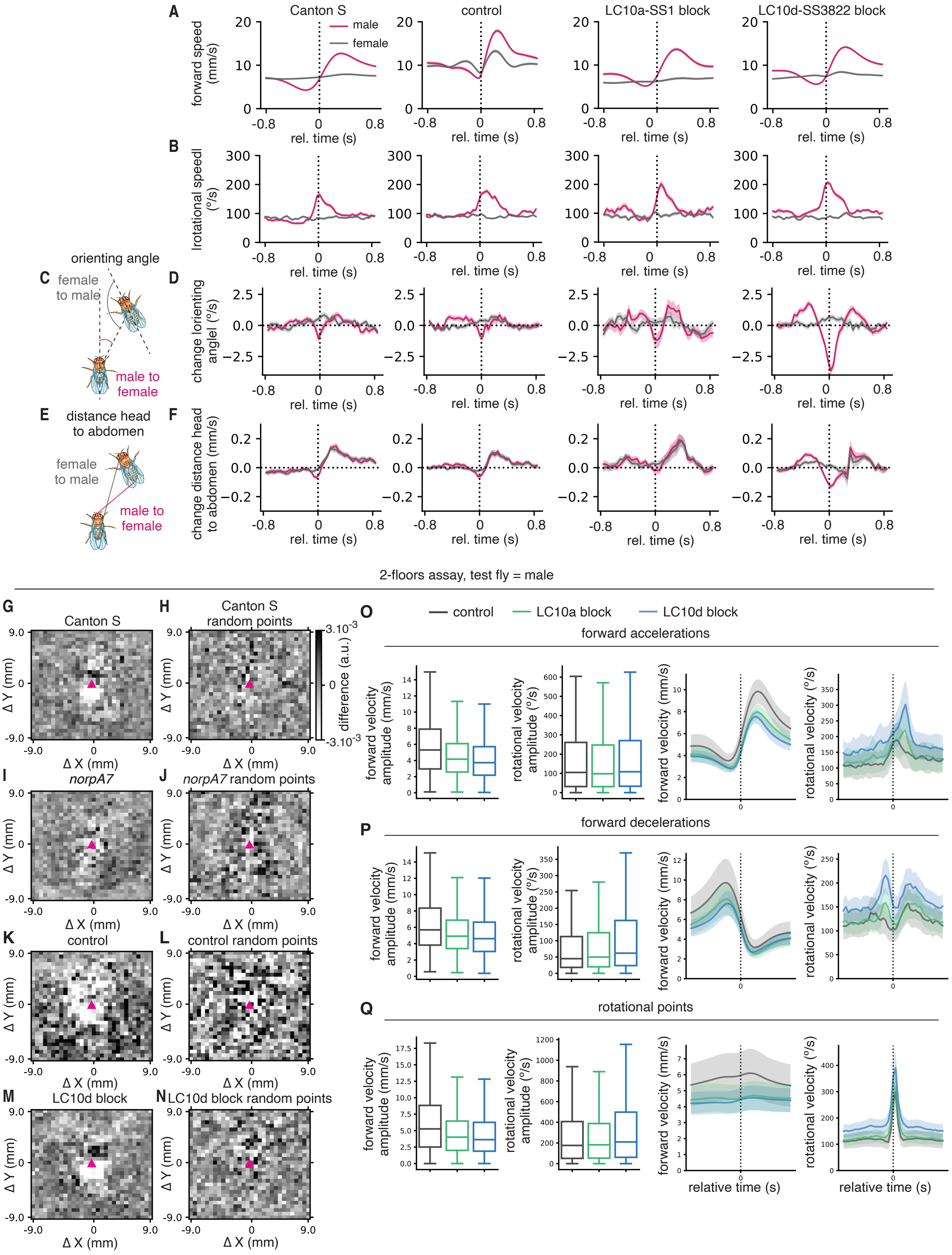

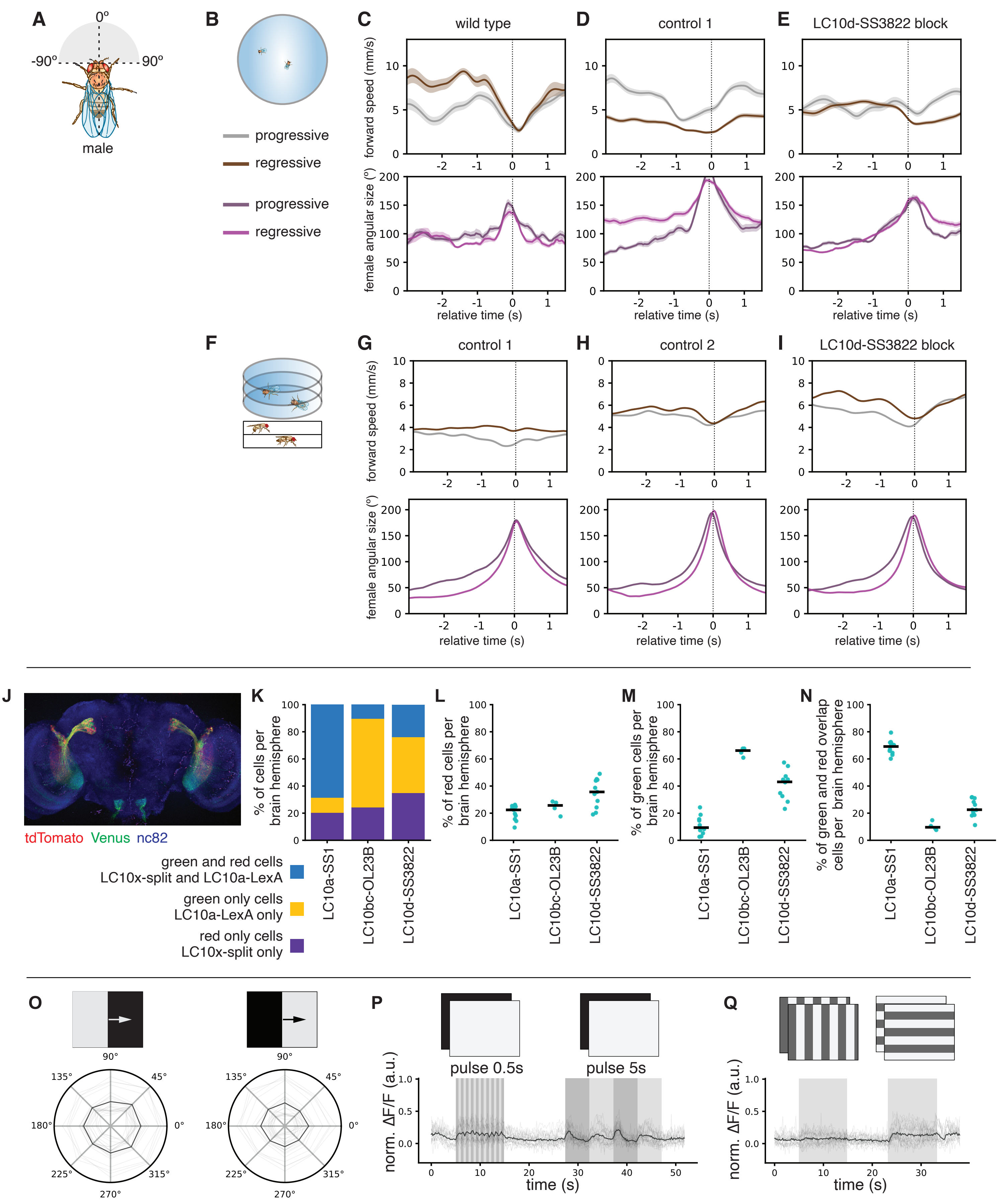

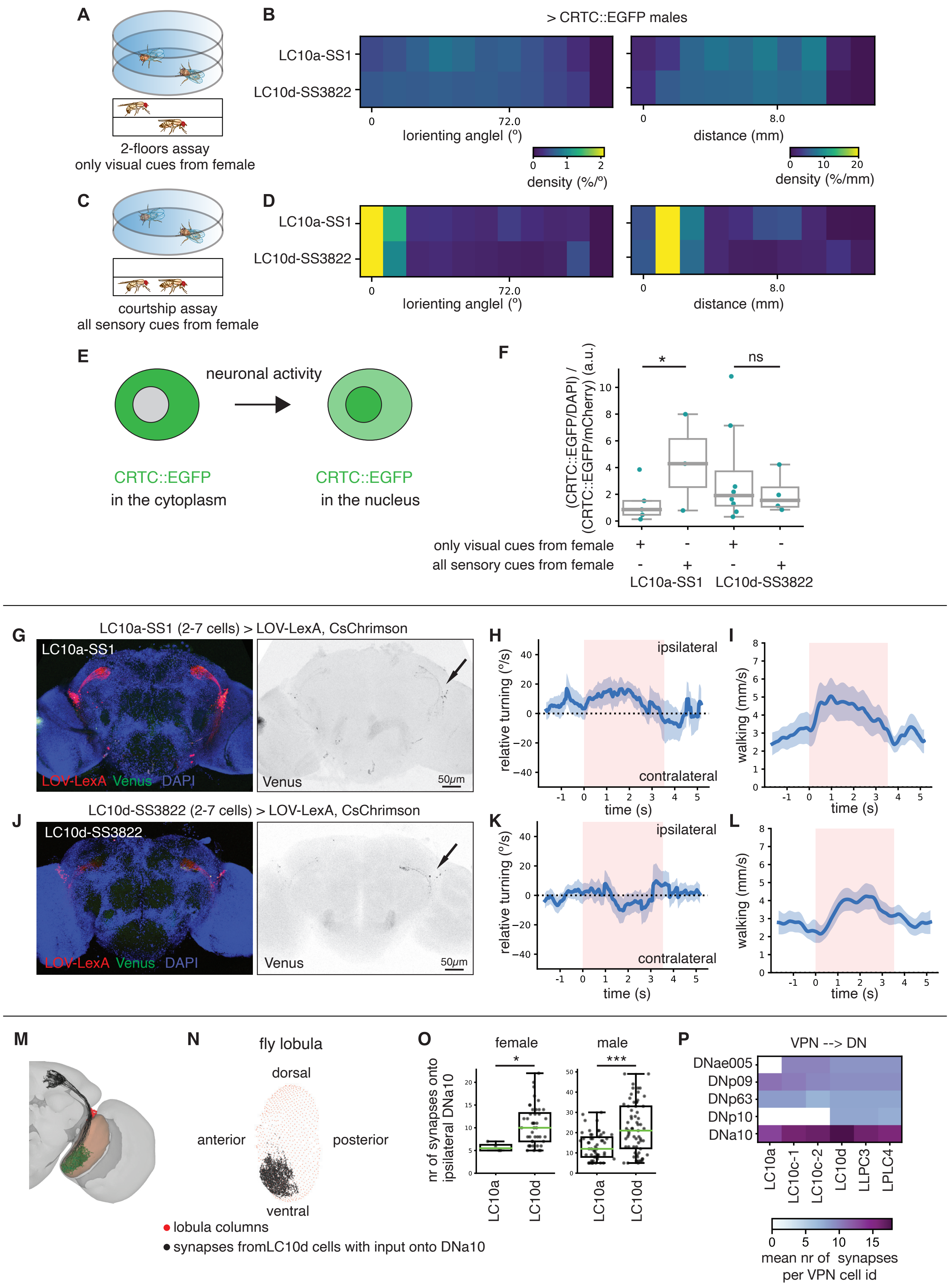

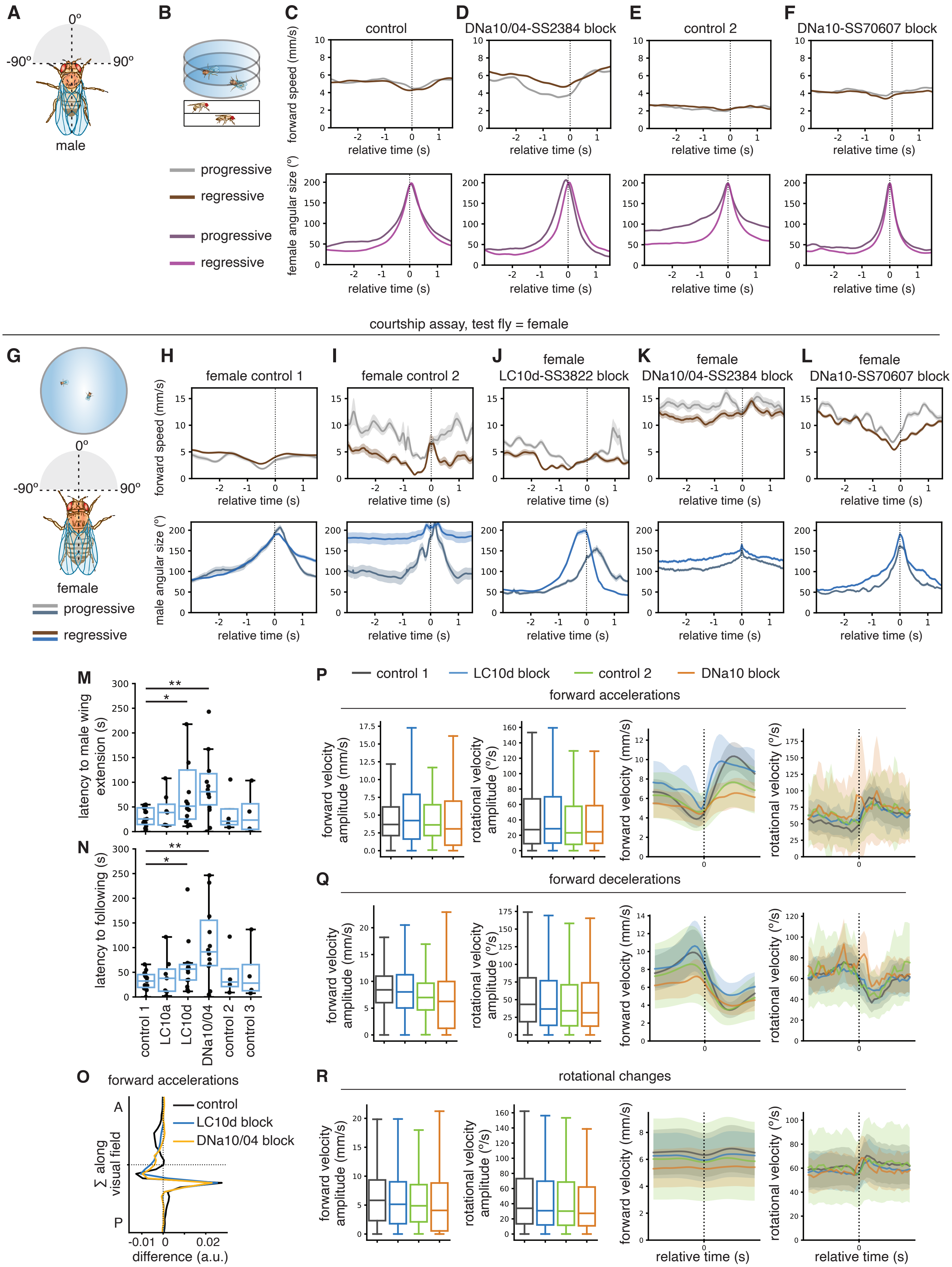
